# Within-host genetic diversity of SARS-CoV-2 across animal species

**DOI:** 10.1101/2024.04.03.587973

**Authors:** Sana Naderi, Selena M. Sagan, B. Jesse Shapiro

## Abstract

Infectious disease transmission to different host species makes eradication very challenging and expands the diversity of evolutionary trajectories taken by the pathogen. Since the beginning of the ongoing COVID-19 pandemic, SARS-CoV-2 has been transmitted from humans to many different animal species, within which viral variants of concern could potentially evolve. Previously, using available whole genome consensus sequences of SARS-CoV-2 from four commonly sampled animals (mink, deer, cat, and dog) we inferred similar numbers of transmission events from humans to each animal species. Using a genome-wide association study (GWAS), we identified 26 single nucleotide variants (SNVs) that tend to occur in deer – more than any other animal – suggesting a high rate of viral adaptation to deer. The reasons for this rapid adaptive evolution remain unclear, but within-host evolution – the ultimate source of the viral diversity that transmits globally – could provide clues. Here we quantify intra-host SARS-CoV-2 genetic diversity across animal species and show that deer harbor more intra-host SNVs (iSNVs) than other animals, providing a larger pool of genetic diversity for natural selection to act upon. Mixed infections involving more than one viral lineage are unlikely to explain the higher diversity within deer. Rather, a combination of higher mutation rates, longer infections, and species-specific selective pressures are likely explanations. Combined with extensive deer-to-deer transmission, the high levels of within-deer viral diversity help explain the apparent rapid adaptation of SARS-CoV-2 to deer.

## Introduction

Viral genetic diversity, at the scale of an outbreak or pandemic, ultimately originates within individual hosts. This is because all mutations that spread between hosts must arise from a population of replicating viruses within an infected individual. Natural selection can act on this genetic diversity both within an individual infection and upon transmission to a new host. In acute infections, selection may have limited time to act or have limited genetic diversity to act upon (Lythgoe et al., 2021). Therefore the strongest selective filter may occur upon transmission to new hosts (Xue et al., 2020, N’Guessan et al., 2023). In some cases, viral lineages that are selectively favored within a host can be transmitted to other hosts (Gonzalez-Reiche et al., 2023). As a result, within-host viral diversity may foreshadow the success of viral mutations and lineages at a global scale (Xue et al., 2017, Harari et al., 2022).

Since the beginning of the COVID-19 pandemic, SARS-CoV-2 has been transmitted from humans to many other animal species, and back to humans. Persistence of the virus in animal reservoirs can open new evolutionary trajectories and allow for the selection of novel mutations. This highlights the need to study within-host diversity of SARS-CoV-2 in animal species, as a complement to recent analyses of consensus-level genomic diversity that ignored within-host diversity (Naderi et al., 2023, Tan et al., 2022, Pickering et al., 2022). In a previous genome-wide association study (GWAS) of consensus sequences, we identified 26 SARS-CoV-2 mutations associated with deer and three other mutations associated with mink, suggesting species-specific adaptations (Naderi et al., 2023). Consensus sequences represent the ‘average’ or ‘majority rule’ genome from a potentially genetically diverse population of viruses within a host. The consensus sequence provides a proxy for the genetic variation most likely to have been transmitted from host to host, while ignoring low-frequency mutations more likely to have arisen within a host (Andersen, Shapiro et al., 2015).

Several studies have quantified and analyzed the level of intra-host diversity of SARS-CoV-2 in humans. Intra-host diversity of SARS-CoV-2 is generally low in acute human infections (Valesano et al., 2021, Lythgoe et al., 2021, Braun and Moreno et al., 2021), but much higher in chronic infections, during which mutations can be selected within a host and potentially transmitted to new hosts (Gonzalez-Reiche et al., 2023). In acute infections, the median number of intrahost single nucleotide variants (iSNVs) at measurable frequency ranges from one to eight (Lythgoe et al., 2021, Valesano et al., 2021), with most samples having fewer than three (Lythgoe et al., 2021). The expected number of nucleotide differences between any two genomes within an average human infection has been estimated to be 0.83 (Tonkin-Hill et al., 2021). A measure of selective constraint at the protein level, the ratio of nonsynonymous to synonymous substitutions rates (dN/dS), was similar between consensus sequences (transmitted between hosts) and moderate frequency iSNVs, suggesting similar levels of selection acting within and between hosts (Tonkin-Hill et al., 2021). By contrast, chronic infections, often in immunocompromised individuals (Avanzato et al., 2020, Choi et al., 2020), allow both the time and modest selective pressure (an immune system that is weak enough to allow prolonged viral replication, but strong enough to impose selection for immune evasion) to favor adaptive evolution (Clark et al., 2021, Choi et al., 2020). Many of the mutations selected within chronic SARS-CoV-2 infections are identical to those present in globally circulating variants of concern, strongly suggesting that successfully transmitting variants might arise from chronic infections (Harari et al., 2022).

While previous studies quantified within-host diversity of SARS-CoV-2 in humans, the diversity within animal hosts has yet to be explored systematically. In this study, we analyzed publicly available short-read sequence data from SARS-CoV-2 sampled from cats, dogs, mink, and deer. For comparison, we matched the animal-derived sequences to an equivalent number of human-derived sequences by their geographical location and time of sample collection. As described below, we found that deer have higher intra-host diversity than other animals, providing a larger pool of genetic diversity upon which natural selection can act.

## Methods

### Data

Raw Illumina sequencing reads from SARS-CoV-2 whole-genomes were downloaded from the NCBI Short Read Archive (National Library of Medicine (US)) for cat, dog, mink, and deer hosts as of September 2022. Samples with incomplete metadata were discarded. The dataset consists of 16 samples from cats, 10 from dogs, 66 from minks, and 81 from deer (58 lymph node tissue samples and 23 nasopharyngeal samples). Most of these sequences came from natural infections, with the exception of six dog samples and 12 cat samples derived from laboratory infection experiments. We searched the SRA database in June 2023 for human-derived short read sequences collected in the same geographical location as each group of animal host sequences, and matched them to animal host sequences based on their month of collection. For each animal, for unique month-location combinations, we sampled human-host sequences from that same location, such that their sample collection date was as close to the animal-host sample as possible. Due to the limited number of sequences available in the SRA database in certain time periods, not all human host matches were in the same month as their respective animal-host sequences. For 65 out of the 173 initial animal samples, human matches fell within the same month, for 45 the human matches were 1-2 months apart, for 46, 3-7 months apart, and for three samples 10 months apart. Among the deer samples, 14 sequences from Ontario had best matches to human-derived samples with incomplete data (only one read from a pair) that were therefore excluded. We proceeded with 60 human-derived samples (similar to the average sample size from other species), for a total of 233 samples across all species. The SRA identifiers for these samples and their respective metadata are listed in **Supplementary Table 1**. The distribution of samples from each BioProject ID is shown in **Supplementary Figure 1**.

### Read mapping and variant calling

Reads in FASTQ format were mapped to the Wuhan-Hu-1 reference genome (MN908947.3) using the GenPipes 4.4.5 pipeline (Bourgey et al., 2019). Within this pipeline, rewards are mapped to reference using BWA software (Li et al., 2009). Primers are trimmed using iVar (Grubaugh et al., 2019). Intrahost single nucleotide variants (iSNVs) are called with Freebayes 1.3.4 (Garrison and Marth et al., 2012), generating Variant Call Format (VCF) files that were used for subsequent analyses. The ploidy is set to haploid by default within the Genpipes pipeline. In all subsequent analyses, only samples with a minimum average depth of 50x, and at least 15,000 nucleotide sites (>50% of the genome) with minimum of 50x coverage were retained. The distribution of the breadth of coverage in the samples included in the analyses are shown in **Supplementary Figure 2**. We also performed a sensitivity analysis considering only samples with >75% of the genome covered at 50X.

### Calculating population genetic metrics

From each VCF file, we calculated the total number of iSNVs in the sample and the nucleotide diversity (the average number of pairwise nucleotide differences, π). To minimize the impact of sequencing errors, nucleotide positions covered by fewer than 50 mapped reads, or with a within-sample minor allele frequency less than 5% were excluded. We calculated π, the average number of pairwise differences, as follows:

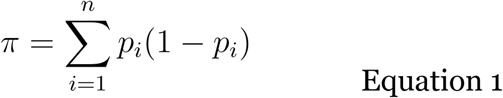

Where *n* is the number of SNVs and *p_i_* is the frequency of an SNV at position *i*, following Tonkin Hill et al., 2021. Corresponding values for each sample are listed in **Supplementary Table 2**.

### Richness and Shannon diversity

To assess within-host lineage diversity, we used the demixing algorithm implemented in Freyja (Karthikeyan et al., 2022) to infer mixtures of SARS-CoV-2 lineages assigned based on the Pango nomenclature, and their relative abundances in each sample, with *richness* defined as the number of unique lineages detected in each individual. Lab-infected samples are known to have been infected experimentally with only one lineage, allowing us to establish a minimum lineage frequency threshold below which only false positive lineage calls are expected. All 18 lab-infected samples were inferred by Freyja to contain only one lineage as long as we included only lineages present at a frequency of 0.025 or more; therefore we used this minimum frequency threshold to remove likely false-positive lineage calls from natural infection samples. We then calculated the Shannon diversity index, *H,* as follows:

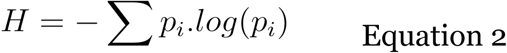

Where *p_i_* is the relative abundance of Pango lineage *i* within a sample. Corresponding values for each sample are listed in **Supplementary Table 2**.

### Determining the representative lineage for samples

For every sample, the most prevalent lineage identified by Freyja after frequency filtering was picked as the representative lineage for that sample. To simplify visualization and modeling the data, we binned groups of closely related lineages together: all B.1.1.x lineages were binned together, all B.1.x lineages were binned together (with the exception of the Delta variant, B.1.617.2), and all other lineages were left unbinned.

### Statistical Modeling

The variables of interest were: host species (with ‘human’ set as the reference level), type of infection (natural vs. lab infection, with natural set as the reference), tissue isolation source (lymph node vs. nasopharyngeal samples, with nasopharyngeal set as reference), sample mean depth of coverage, sample breadth of coverage (percentage of sites across the genome with at least 50x coverage), and assay type (whole genome shotgun vs. amplicon sequencing). We assumed a Poisson distribution for the number of iSNVs and richness, and modeled π and Shannon diversity using linear models. Since lymph node tissue was sampled exclusively from deer, with the exception of **model S1**, we chose to construct our models on nasopharyngeal samples only to avoid complications from the collinearity of species and tissue type, and separately tested for tissue effects using data from deer only.

### Generalized Linear Mixed Models

All Generalized Linear Mixed Models (GLMMs) were fitted using the *glmer* function in the *lme4* package (*1.1.34,* Bates D et al., 2015) in R (*4.3.1*, R Core Team, 2023). The number of maximum iterations for all GLMMs was set to 2 x 10^5^, and the optimizer “bobyqa” was used. The number of iSNVs and richness were modeled using GLMMs, and a Poisson distribution was assumed (*family* argument set to “poisson”).

### Linear mixed models

All Linear Mixed Models (LMMs) were fitted using the lmer() function from the R (*4.3.1, R Core Team, 2023*) *lmerTest* package (*3.1.3.* Kuznetsova ET AL., 2017,). π and Shannon diversity were modeled using LMMs.

### Generalized Linear Models

The Poisson Generalized Linear Model described in the text was run using the *glm()* function in the stats package (*4.3.1,* R Core Team, 2023) in R (*4.3.1*, R Core Team, 2023). The *family* argument was set to “poisson” with a “log” link.

### Multicollinearity analysis and adjusted Variant Inflation Factors

Multicollinearity was assessed on the fitted models using the adjusted Generalized Variance Inflation Factor (GVIF) calculated in terms of the correlation matrix of the regression coefficients (Fox et al., 1998) by the *vif()* function from the CAR package (*3.1.2*, Fox et al., 2019) in R (*4.3.1*, R Core Team, 2023). As a rule of thumb, an adjusted GVIF of 2.23 (square root of 5, equivalent to the threshold commonly used in VIF analyses for predictors with one degree of freedom) and higher was used to consider multicollinearity as ‘high’, adjusted GVIF values above 1.75 were considered moderate and were further investigated, and adjusted GVIF values below 1.75 were considered ‘low’.

#### Description of fitted models

**Model 1.** Poisson GLMM for the number of iSNVs as the response variable, fitted on nasopharyngeal samples. Fixed effects in this model were species, type of infection, depth and breadth of coverage, and assay type. BioProject ID was a random effect to control for project-specific differences in sample handling (*e.g.* cross-contamination) or sequencing quality. This model had no singularity or multicollinearity issues, and all adjusted GVIF values were low (< 1.75).

**Model 2.** LMM for π, fitted on nasopharyngeal samples. Fixed effects in this model were species, type of infection, depth and breadth of coverage, and assay type. BioProject ID was a random effect. This model had no singularity or multicollinearity issues, and all adjusted GVIF values were low (< 1.75).

**Model 3.** Poisson GLMM for lineage richness, fitted on nasopharyngeal samples. Initially, host species, type of infection, breadth and depth of coverage and assay type were set to fixed effects, and BioProject ID was a random effect. The model resulted in a near singular fit and though the adjusted GVIF values were all below the cutoff, they were moderate (1.90 for type of infection, due to collinearity of assay type with type of infection in this model). Removing assay type from the fixed effects reduced the adjusted GVIF values (all adjusted GVIF values below 1.75) and resolved the singularity issue of the model fit.

**Model 4.** LMM for Shannon diversity, fitted on nasopharyngeal samples. Fixed effects in this model were species, type of infection, depth and breadth of coverage, and assay type. BioProject ID was a random effect. This model had no singularity or multicollinearity issues, and all adjusted Generalized Variance Inflation factors were low (< 1.75).

**Model 5**. Poisson GLMM for the number of repeated iSNVs in humans, deer, and mink. The outcome in this model was the percentage of samples with repeated iSNVs (at least two in a given species), and the predictors were species and mutation type.

**Model 6**. Poisson GLM for the number of repeated iSNVs across deer samples. We only included iSNVs that occur in two or more samples, and only those in Open Reading Frames (ORFs). When an iSNV occurs in a region of overlapping ORFs, it is counted twice. The outcome in this model was the percentage of samples in which the same iSNV appeared, and the predictor was type of mutation (synonymous or non-synonymous).

**Model S1.** Poisson GLMM for the number of iSNVs, on all nasopharyngeal and lymph node tissue samples. Fixed effects in this model were species, type of infection, tissue type, depth and breadth of coverage, and assay type. BioProject ID was a random effect. This model had no singularity or multicollinearity issues, and all adjusted GVIF values were low (< 1.75).

**Model S2.** Poisson GLMM for the number of iSNVs, on deer samples only. Fixed effects in this model were tissue type, depth and breadth of coverage (type of infection, species, and assay type are invariable in the deer samples and thus are no longer predictors). BioProject ID was a random effect in the model. This model had no singularity or multicollinearity issues, and all adjusted GVIF values were low (< 1.75).

**Model S3.** LMM for π, on deer samples only. Fixed effects in this model were tissue type, depth and breadth of coverage (type of infection, species, and assay type are invariable in the deer samples and thus are no longer predictors). BioProject ID was a random effect in the model. This model had no singularity or multicollinearity issues, and all adjusted GVIF values were low (< 1.75).

**Model S4.** Poisson GLM for richness, on deer samples only. We initially aimed to build the model with a GLMM, but any GLMM combination with tissue type and BioProject ID resulted in a singular fit, despite adjusted GVIF values being low. We believe this is due to the overcomplexity of the model relative to the sample size. With tissue type being the primary effect of interest, we resorted to a Poisson GLM instead, where tissue type, depth and breadth of coverage were predictors. This model had no singularity or multicollinearity issues, and all adjusted GVIF values were low (< 1.75). We interpret the results of this model with caution as we were not able to control for the effects of BioProject ID as a random effect.

**Model S5.** LMM for Shannon diversity, on deer samples only. The initial model with tissue type, breadth and depth of coverage as fixed effects and BioProject ID as a random effect had a singular fit, despite low adjusted Generalized Variance Inflation Factor values. The only model combination with both tissue type and BioProject ID without a singularity issue, was when breadth of coverage was excluded from the model. The alternative option being a simple Linear Model, we preferred excluding breadth of coverage to not controlling for BioProject, and constructed the model with tissue type and depth of coverage as fixed effects, and BioProject ID as a random effect. This model had no singularity or multicollinearity issues, and all adjusted GVIF values were low (< 1.75).

**Model S6.** GLMM for the number of iSNVs, on nasopharyngeal samples only and controlling for lineage effects. Fixed effects in this model were species, type of infection, breadth and depth of coverage, and assay type. Random effects in this model were BioProject ID and representative lineage. This model had no singularity or multicollinearity issues, and all adjusted GVIF values were low (< 1.75).

**pN/pS.** We used the genetic distance-based method (Nei, M., and T. Gojobori. 1986) to calculate the ratio of nonsynonymous to synonymous within-host polymorphism rates within each sample (thus the name, pN/pS). The difference between our measure of pN/pS (based on iSNVs) and the between-host dN/dS measure is that the reference allele at each segregating site was set to the major allele, under the assumption that observed minor alleles likely appeared within the sample. Other than rare cases of mixed infections (detected by Freyja above), all sequences within a sample share a common ancestor at the time of infection (Lythgoe et al., 2021). Over this relatively short evolutionary time, we considered it rare for a codon to contain more than one mutation event. For each sample, we calculated pN/pS as follows:

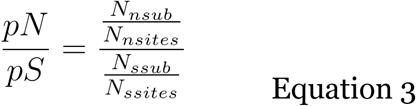

Where *N_nsub_* and *N_ssub_* are the number of observed nonsynonymous and synonymous iSNVs within the sample, *N_nsites_* and *N_ssites_* are the number of nonsynonymous and synonymous sites, respectively. We only computed the pN/pS ratios in samples with one or more synonymous iSNVs (pS > 0).

pN/pS was calculated both genome-wide, and for the S gene alone. We used the non-parametric Wilcoxon rank sum test to compare pN/pS values genome-wide and in the S gene for each animal species versus humans. The test was run using the *wilcox.test* function in the R stats package (R Core Team, 2023), and significance was tested at a 0.05 level.

### Repeated iSNVs across samples from the same species

To identify individual sites under positive selection, we counted the number of times each of the observed iSNVs repeatedly at the same nucleotide site across the samples of each species. The same filtering criteria as described before were used to exclude mutations potentially arising from sequencing errors. We defined a ‘repeated iSNV’ as a substitution that appears in two or more samples from the same species. For mutations in overlapping ORFs, we counted the substitution twice, and determined its type (synonymous or nonsynonymous) relative to each of the ORFs it belongs to. Intergenic mutations were excluded from the analysis.

### Detecting animal-specific mutations within hosts

Previously, we used a genome-wide association study (GWAS) to identify mutations in the SARS-CoV-2 genome associated with different animal species compared to humans. For each of the previously identified 26 deer-associated and three mink-associated mutations (Naderi et al., 2023) we counted the number of times they appear as “fixed” or “ambiguous” (iSNVs) in the “consensus tag” column of the VCF files generated from deer and mink samples. We then counted the same quantities for every site in the SARS-CoV-2 that was not associated with either of these animals (non-GWAS hits). The categorization was done based on the Consensus Tag for each mutation in the VCF file generated by Freebayes (Garrison and Marth et al., 2012). The counts for each animal were tallied to compare GWAS and non-GWAS sites. The detailed breakdown of the counts for each GWAS hit is listed in **Supplementary Table 3**.

## Results

### Deer contain relatively high within-host SARS-CoV-2 diversity

To compare within-host diversity of SARS-CoV-2 across animal species, we first counted the number of iSNVs within samples taken from cats, dogs, deer, mink, or humans. The median number of two iSNVs within human samples in our dataset was similar to previous estimates from acute human infections (Lythgoe et al., 2021) and also similar to mink (**Figure 1a**). By contrast, deer nasopharyngeal samples had a higher median of four iSNVs per sample compared to equivalent human samples (**Figure 1a**). To test the statistical significance of these differences while controlling for sequencing coverage and covariates such as type of infection, we used generalized linear mixed models (**Methods; GLMMs**). Our goal was to determine how the number of iSNVs varied across animal species compared to humans as a reference. Due to the collinearity of species and tissue type (all lymph node tissue samples were isolated from deer), we chose to construct our models on nasopharyngeal samples only. We fitted a GLMM with host species, type of infection, breadth and depth of coverage, and assay type as fixed effects, and BioProject ID as a random effect to control for project-specific technical differences (**Methods, Model 1**). Host species and breadth of coverage were significant predictors in this model (**Table 1**). Deer had a significantly higher number of iSNVs compared to humans, with a rate ratio 2.71, indicating that the number of iSNVs is 171% higher in deer than human nasopharyngeal samples. To more conservatively exclude potentially lower quality samples, we repeated Model 1 while requiring that 75% rather than 50% of the genome be covered at 50X, which excluded a few outlying samples (**Supplementary Figure 2**). With this more conservative filtering, deer still had significantly higher numbers of iSNVs than humans (rate ratio = 2.64, *p* =0.02). An equivalent full GLMM for the number of iSNVs including lymph node tissue samples along with the nasopharyngeal samples yielded similar results, with deer having significantly more iSNVs than humans (**Methods, Model S1**, **Supplementary Table 4**).

**Figure 1.**
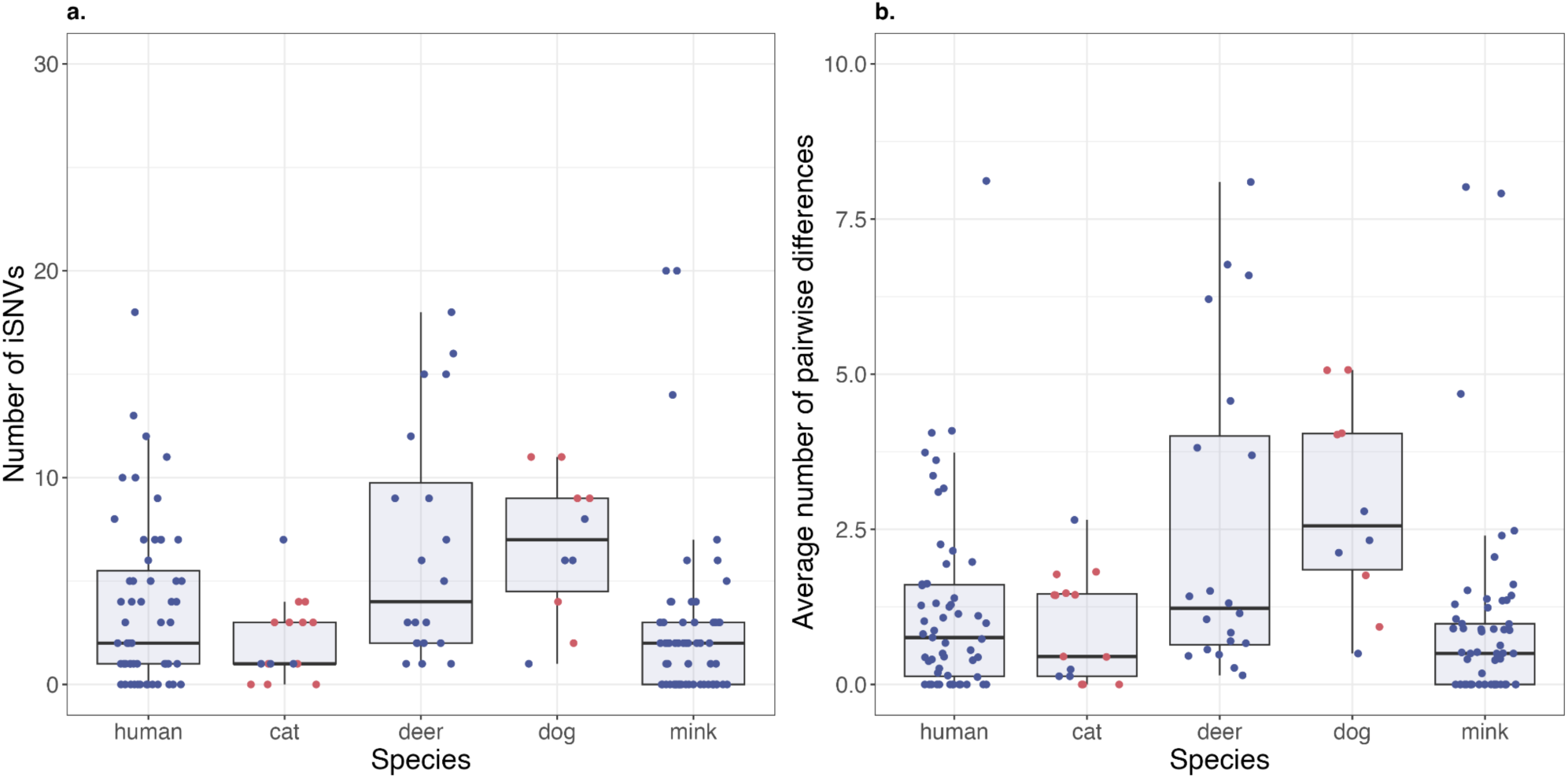
Higher within-host viral diversity in deer than humans. Boxplots display (a) the number of iSNVs and, (b) the average number of pairwise differences (π) across animal species on nasopharyngeal samples only. Raw data points are shown with random horizontal jitter. Blue points are from natural infections and pink points are from experimental lab infections. The boxes display the interquartile range, with the midline marking the median, and whiskers extending up to 1.5x of the interquartile range, or data extremums. Datapoints beyond whiskers are outliers according to the interquartile criterion.

**Table 1:** Table of coefficients for **Model 1**. Poisson GLMM, fitted on nasopharyngeal samples only with the number of iSNVs as the response variable. Significant effects (*P* < 0.05) are highlighted in pink. See **Model S1 (Supplementary** Table 4) for the full model including lymph node tissue samples.

| Predictors (Fixed effects) | Estimate | Rate ratio | P-value |
| --- | --- | --- | --- |
| **Species (Reference: human)** |  |  |  |
| cat | NA | NA | p>0.05 |
| dog | NA | NA | p>0.05 |
| mink | NA | NA | p>0.05 |
| deer | 0.9964 | 2.7085 | 0.0355 |
| **Type of infection (Reference: Natural infection)** |  |  |  |
| Lab infected | NA | NA | p>0.05 |
| **Breadth of coverage** | -1.9331 | 0.1447 | 1.6170E-05 |
| **Depth of coverage** | NA | NA | p>0.05 |
| **Assay type** | NA | NA | p>0.05 |

Lymph node tissue was only sampled from deer, and had a higher median number of iSNVs than nasopharyngeal deer samples (median of 4 and 21 iSNVs for nasopharyngeal and lymph node tissue samples, respectively, **Supplementary Figure 3a**). However, in the full GLMM model (**Model S1**) where lymph node tissue samples were also included (**Supplementary Table 4**), tissue isolation source was not a significant effect. Moreover, a GLMM fitted on deer samples alone did not reveal a significant difference in the two tissue types. This deer-specific model had tissue isolation source, depth and breadth of coverage as fixed effects, and BioProject ID as a random effect (**Model S2, Supplementary Table 5**). The results of Models S1 and S2 demonstrate that the difference between the number of iSNVs in the two tissue types in deer (**Supplementary figure 3a**) cannot be separated from project-specific technical differences.

Breadth of coverage is a significant effect in Model 1, with a rate ratio of 0.14, indicating that samples with lower breadth of coverage tend to contain a higher number of iSNVs (**Table 1**). This could be a sequencing artifact, with more poorly covered sequences prone to having a higher number of false-positive iSNVs called. Importantly, deer still have more iSNVs than humans even after accounting for breadth of coverage in the model. We also note that contrary to the expectation that lab infections are of short duration and might have less time to accumulate iSNVs, we observe a higher number of iSNVs in lab-infected samples (median of 1 and 3 iSNVs for natural and lab cat infections respectively, and median of 6 and 9 iSNVs for natural and lab dog infections respectively). This effect is equalized once BioProject ID is controlled for, and type of infection (lab vs. natural) was not statistically significant in the model (**Table 1**).

Next, we compared π, the average number of pairwise nucleotide differences within a sample. Unlike the number of iSNVs, which treats variants at all frequencies equally, π is maximized when variants approach intermediate frequencies and is less sensitive to low-frequency variants. We observe that deer have higher median values of π compared to humans (median of 1.23 for deer, and 0.57 for humans, **Figure 1b**). While dogs also had higher median π values (median of 2.44 for dog, **Figure 1b**), but this result relied on a relatively small sample of dogs.

We sought to statistically test the differences in π across host species using a Linear Mixed Model (LMM). In the LMM for π (**Model 2**), species, type of infection, depth and breadth of coverage and assay type were fixed effects, and BioProject ID was a random effect. The model was fitted on nasopharyngeal samples only. Deer was significant in this model with an estimated slope of 1.54 (p = 0.037, **Supplementary table 6**), indicating that if all other variables are held constant, deer have a higher π than humans, with an expected difference of 1.54. This is consistent with the distribution of minor allele frequencies across species, with a shift toward more intermediate frequencies in deer (**Supplementary Figure 4**). With other covariates such as assay type and infection type controlled for, π within dogs was not significantly different than in humans **(Supplementary Table 6)**.

We next fitted a deer-specific LMM for π to assess the effect of tissue isolation source, while controlling for sequencing quality. In this LMM, tissue isolation source, depth and breadth of coverage were fixed effects, and BioProject ID was a random effect (**Model S3, Methods**). Despite having a higher median π, we found that lymph node tissue samples were not statistically different from nasopharyngeal samples (**Supplementary Figure 3b, Supplementary table 7**). The observed difference between lymph and nasopharyngeal deer samples is likely due to sequencing quality, and is controlled for by project-specific variabilities. This is consistent with the analysis of iSNV number, in which tissue-specific differences could not be separated by technical confounders (**Model S2, Supplementary table 5**).

### Mixed infections are rarely detected

Within-host genetic diversity can emerge via *de novo* mutations occurring during infection or from mixed infections of lineages that diverged before the animal was infected. To determine whether mixed infections could have contributed to the observed within-host diversity described above, we inferred the presence and relative abundance of distinct lineages of SARS-CoV-2 within each sample (**Methods**).

Despite eliminating low frequency lineages whose frequency fell below our minimum threshold of 0.025 (based on lab-derived samples, infected with a single isogenic viral stock; Methods), multiple viral lineages were inferred in 68 out of the 233 samples. The lineages inferred within a sample were almost always closely related, suggesting false-positive coinfections that could plausibly have arisen instead by *de novo* mutation or sequencing errors. Of the 68 samples inferred to contain more than one distinct lineage, only one (from a human sample) contained two distinct WHO-defined variants of concern, Alpha and Gamma (**Supplementary Table 8**). The one human-host sample with 22 inferred lineages only contained closely related (B.1.1.x) lineages, which could be false positives. Seven other samples contained mixtures of Delta and multiple closely-related B.1.x lineages. We consider these eight samples to be plausible mixed infections due to the inferred presence of at least two distantly related lineages. In the remaining 60 samples, all identified lineages were closely-related B.1.x descendants (**Supplementary Table 8**), which we consider dubious mixed infections that are difficult to separate from *de novo* mutation or technical artifacts.

Considering nasopharyngeal samples alone, we find the same median number of lineages for deer and humans (median richness of one lineage per sample in both humans and deer nasopharyngeal samples). After controlling for effects of coverage and infection type, the only species with a significantly different richness than humans was mink (**Table 2**). The GLMM fit for lineage richness included species, type of infection, breadth and depth of coverage as fixed effects, and BioProject as a random effect (**Model 3, Table 2**). Assay type was removed from this model due to multicollinearity issues (**Methods**). In this model, mink had a rate ratio of 0.48, indicating that they tend to harbor 52% fewer lineages per sample than humans. Breadth of coverage was a significant effect with a rate ratio of 0.073, indicating that samples with a lower breadth of coverage tend to have a higher number of inferred lineages, highlighting the importance of controlling for coverage effects in these analyses. Together, these results show that the higher within-host diversity in deer compared to humans (**Figure 1**, **Table 1**) is not easily explained by co-infections by different lineages (**Figure 2a**, **Table 2**).

**Figure 2.**
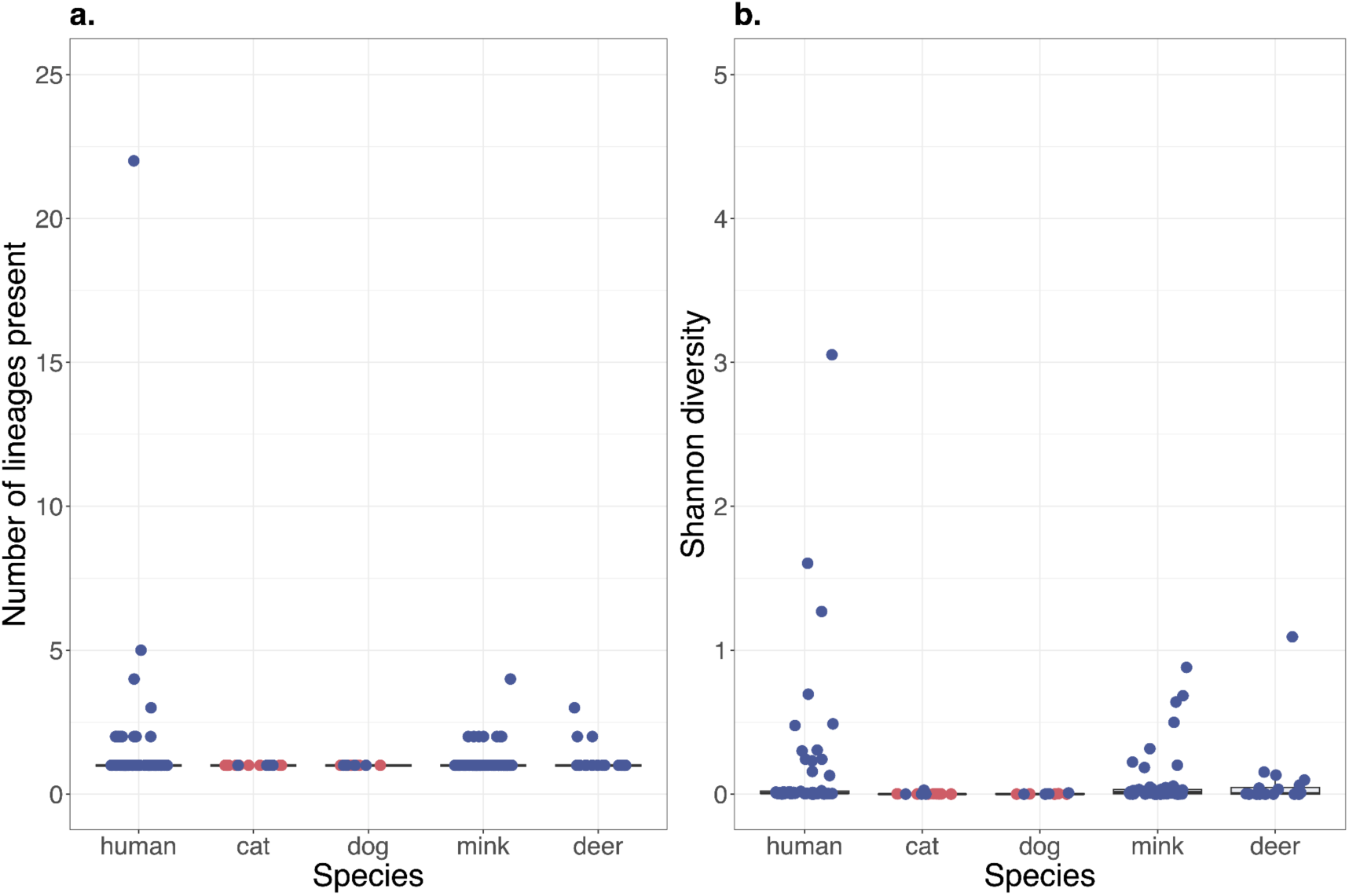
Multiple-lineage infections are rare across species. Boxplots display (a) the number of inferred lineages present and (b) the Shannon diversity index across species, both on nasopharyngeal samples only. Blue points are from natural infections and experimental lab infections in pink. The boxes display the interquartile range, with the midline marking the median, and whiskers extending up to 1.5x of the interquartile range, or data extremums. Datapoints beyond whiskers are outliers according to the interquartile criterion.

**Table 2:**
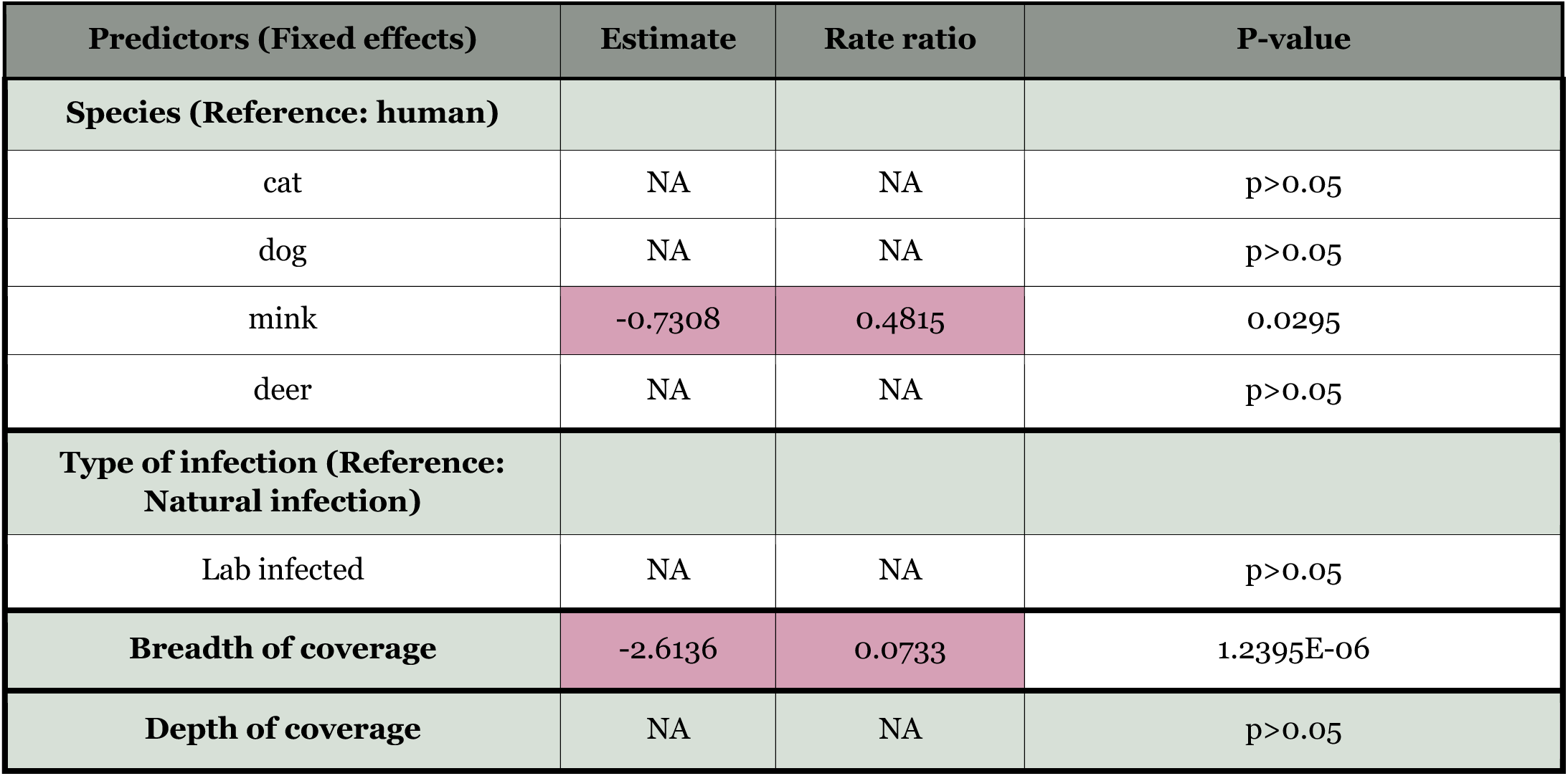
Results of the Poisson GLMM for lineage richness, excluding lymph node tissue samples (**Model 3**). Significant effects (*P* < 0.05) are highlighted in pink.

We further considered the Shannon diversity index, a metric that accounts for both lineage number and their frequencies within a host. The Shannon diversity values for cat and dog were the lowest (median of 0.0010 and 0.0009 respectively, **Figure 2b**), compared with mink, deer, and humans (medians of 0.0134, 0.0062, and 0.0069, respectively, **Figure 2b**). To statistically test these differences, we applied a LMM to nasopharyngeal samples, with species, type of infection, depth and breadth of coverage, and assay type as fixed effects, and Bioproject ID as a random effect (**Model 4**). According to this model, cats and dogs had significantly lower Shannon diversity than humans (**Supplementary Table 9**). This result is expected as mixed infections were never inferred in cats and dogs, which only contain one lineage per sample (**Figure 2a**). However, deer and humans are not significantly different, reinforcing that differences in lineage diversity do not explain differences in their within-host diversity.

In deer, we observed a higher median richness in lymph node tissue samples (median of 3 lineages) than in nasopharyngeal samples (median of 1 lineage, **Supplementary Figure 5a**). To determine the statistical significance of this difference, while controlling for other covariates, we fitted a deer-specific Poisson GLM (Methods, **Model S4**). In this model, tissue isolation source, depth and breadth of coverage were included. Lymph node tissue was a significant effect (p < 0.001, **Supplementary Table 10**), with a rate ratio of 2.77, indicating higher richness in deer lymph node tissue compared with nasopharyngeal. We interpret this result with caution, as for this model, BioProject ID was excluded as a random effect due to singularity issues (**Methods**). The model is therefore not controlled for variability among projects. Furthermore, in a LMM for Shannon diversity, which did include BioProject ID as a random effect (**Model S5**), there were no significant differences between nasopharyngeal and lymph node tissue (**Supplementary Table 11**) despite an observed higher median (**Supplementary Figure 5b**). This supports the conclusion that the higher median Shannon diversity observed in lymph node samples (median 0.006 and 0.820 in nasopharyngeal and lymph samples, respectively) can be explained by project-specific technical differences.

### Lineage-specific differences do not explain differences in within-host diversity across animals

Having established that most samples are likely infected by a single viral lineage which then diversifies by mutation within the host, we considered that lineages could differ in their mutation rate, potentially explaining some of the variation across species. Deer, mink, and human infections are all dominated by lineages B.1.x and B.1.1.x (**Supplementary Figure 6**), suggesting that differences in diversity among these host species are unlikely driven by lineage-specific effects. In contrast, cats and dogs are mostly infected by lineage A, which is not represented in our human samples. In theory, any differences in diversity between humans and cats or dogs could be explained by lineages with different mutation rates. However, in practice, this is not a major consideration since we did not detect any such differences between these animals.

To statistically quantify the effect of lineage in comparisons of within-host diversity across animal species, we added “representative lineage” as a random effect to the GLMM of Model 1 (**Model S6**). The representative lineage is effectively the consensus sequence within a host, and is the likely ancestral genomic background upon which within-host mutations occurred (**Methods**). This model had species, type of infection, depth and breadth of coverage, and assay type as fixed effects, BioProject ID and representative lineage as random effects, and was fitted on nasopharyngeal samples only. In this model, deer have significantly more iSNVs than humans (p = 0.019) with a rate ratio of 3.32, even after controlling for potential lineage effects (**Supplementary Table 12**). We therefore conclude that the higher rates of iSNVs in deer is not likely due to a distinct subset of viral lineages infecting deer compared to humans.

### Testing for selection within-hosts

To provide a coarse-grained measure of natural selection acting on within-host diversity, we compared the ratio of nonsynonymous and synonymous polymorphism rates within each host (pN/pS). In the absence of purifying selection against nonsynonymous mutations, which tend to be deleterious, we expect a pN/pS ratio close to one. By contrast, pN/pS < 1 points to stronger purifying selection against non-synonymous mutations, while pN/pS > 1 is interpreted as positive selection for adaptive protein changes. To avoid division by zero, we excluded samples without any observed synonymous mutations. We observe genome-wide pN/pS < 1 in all species, with medians of 0.56 in cat, dog, and deer, 0.44 in mink, and 0.41 in humans (**Supplementary Figure 7a**). None of the animals have pN/pS significantly different from humans (Wilcoxon rank-sum test, *P* > 0.05 after Bonferroni correction). Most of the outlying deer samples with pN/pS > 1 are from lymph node tissue. This could potentially indicate relaxed purifying selection or strong positive selection in this tissue; however, as mentioned above, these tissue effects are confounded by BioProject ID and thus should be interpreted with caution.

The Spike protein is a major target of the immune system and might be under distinct, and possibly species-specific, selective pressure compared to other genes. We found the range of median pN/pS values in Spike are lower than the genome-wide values with a median of 0.14, 0.28, and 0.43 for human, deer, and mink, respectively (**supplementary figure 7b**). The Spike pN/pS values in mink or deer were not significantly different than humans (Wilcoxon rank-sum test, *P* > 0.05 after Bonferroni correction). Dogs and cats were not tested due to the small number of iSNVs resulting in zero synonymous mutations. Due to the exclusion of samples without synonymous mutations, our sample size is quite small and our power of statistical inference is limited. Nevertheless, these results are consistent with the predominance of purifying selection across the genome within every species, but this does not exclude the possibility of individual nucleotide sites under positive or relaxed purifying selection.

To identify individual sites under positive selection within hosts, we looked for repeated iSNVs across samples of the same species. If a mutation is advantageous in a given host, we expect it to appear repeatedly (twice or more) in independent samples from the same host species. Considering nasopharyngeal samples only, we scaled the repeated iSNV counts to the sample size for each species (excluding dogs and cats due to low counts) and plotted these scaled counts across the genome (**Figure 3**). As an empirical null against which to detect sites with unexpectedly high rates of repeated mutation, we calculated a 95% confidence interval (CI) of repeated synonymous iSNVs (blue shaded areas in **Figure 3**). Visually, it is clear that deer have higher rates of repeated mutations than mink or humans, and that nonsynonymous iSNVs tend to be highly repeated. These effects are significant in a Poisson GLM (**Model 5**). According to this model, deer have significantly higher rates of repeated mutation than humans (*P* < 0.001, rate ratio 1.97), while in mink have lower rates (*P* < 0.001, rate ratio 0.57). Considering only deer samples (**Model 6**), we find that nonsynonymous mutations have higher repeat percentages than synonymous mutations (*P* < 0.05, rate ratio 1.42). The elevated rate of repeated mutation in deer could be explained by higher mutation rates in deer overall (McBride et al., 2023), but specific iSNVs (particular nonsynonymous iSNVs) occurring in more samples than expected based on the 95% CI for synonymous iSNVs are difficult to explain without invoking site-specific and species-specific variation in selective pressures. The results are thus consistent with within-host positive selection on specific nonsynonymous sites, with the precise sites under selection varying by host species (**Supplementary table 13**). When deer lymph node tissue samples are included, we observe similar trends of higher repeat percentages in deer than in humans (*P* < 0.05, rate ratio 1.2), and higher counts rates of non-synonymous repeated iSNVs (P < 0.001, rate ratio 0.35). We also observe an excess of repeated mutations in deer compared to humans and mink, likely driven by lymph node tissue samples. For example, there is an apparent hotspot of repeated nonsynonymous in the nucleocapsid (N) gene in deer, but not in other species (**Supplementary figure 8, supplementary table 14**).

**Figure 3.**
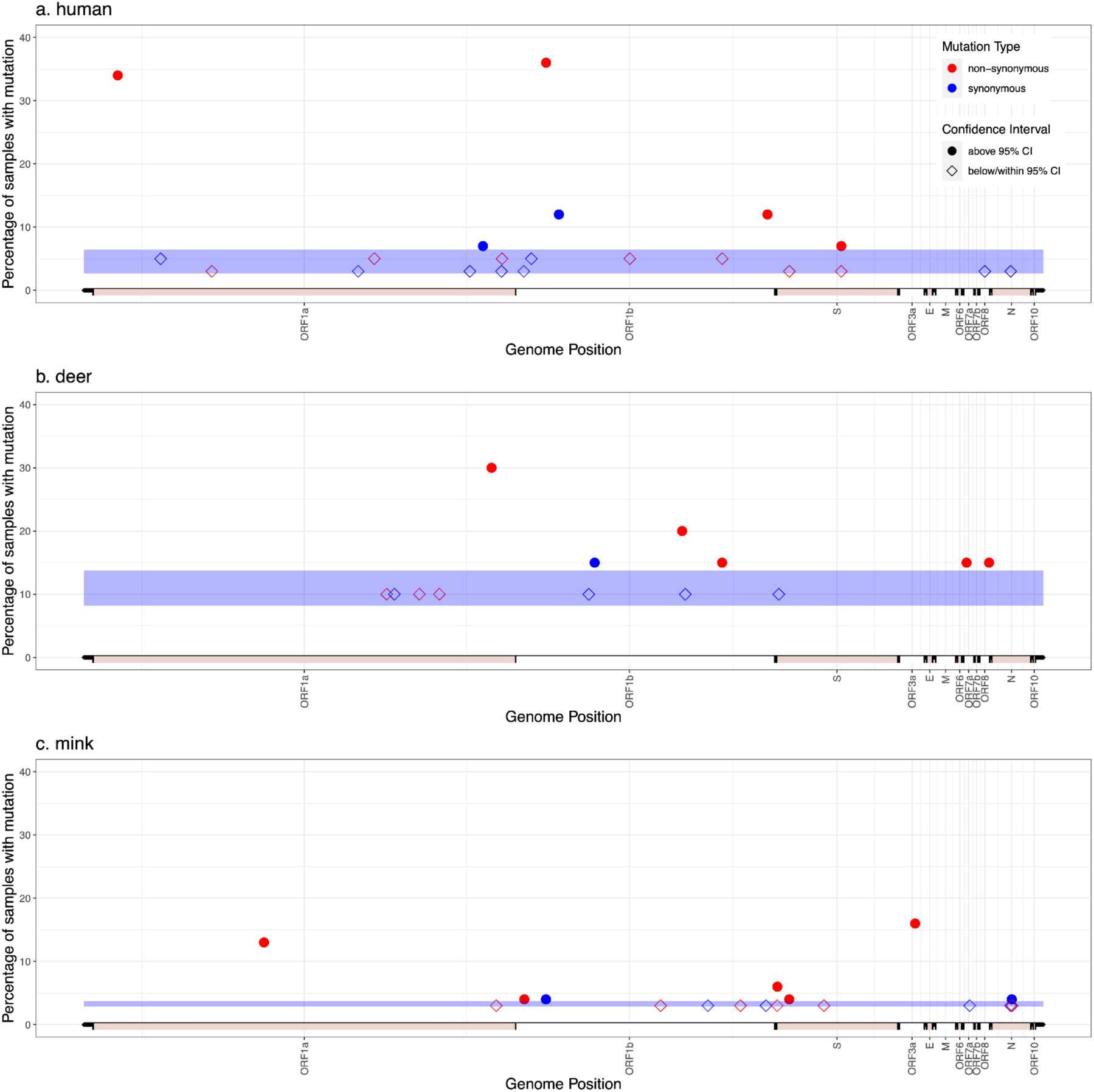
Higher number of repeated iSNVs in deer than human and mink. The distribution of repeated iSNVs across the samples of each species is shown. The y-axis shows the position on the genome, and the y-axis shows the percentage of that species samples harboring a specific iSNV. Synonymous and non-synonymous mutations are in blue and red, respectively. A 95% confidence interval (CI) was calculated using repeated synonymous iSNVs in each species (shaded blue region). iSNVs with repeat percentages above the 95% CI are shown as filled circles, and those within or below the 95% CI are shown as empty diamonds. Only nasopharyngeal samples are included in this analysis.

Some of the repeated iSNVs may provide a selective advantage within a host, but they may or may not be adaptive or transmissible between hosts. If they are involved in longer-term species-specific adaptation, they would be expected to be found as hits in our previous GWAS of consensus sequences (Naderi et al., 2023). We find that some repeated iSNVs are also found as between-host GWAS hits, but many are not. This suggests that some iSNVs are adaptive both within and between hosts, but many are only adaptive within hosts. In mink samples, we observe one of the three nonsynonymous GWAS hits as a repeated iSNV, but it is only repeated in two samples (**Supplementary table 13**). In deer nasopharyngeal samples, we observe one repeated iSNV out of the 21 deer GWAS hits in open reading frames. This iSNV is synonymous, repeated in two deer samples. With lymph node samples included, we observed 16 out of the 21 GWAS hits as repeated iSNVs deer, nine of which are above the 95% CI (**supplementary table 14**).

To more formally test if iSNVs are more likely to be involved in longer-term (between-host) species-specific adaptation than expected by chance, we counted the number of repeated iSNVs at nucleotide positions identified as GWAS hits compared to other sites in the genome. We found that deer-associated GWAS hits are more likely to be found as polymorphic (iSNVs) within deer samples compared to SNVs that were not GWAS hits, and compared to mink-associated GWAS hits, which are not expected to be adaptive in deer (**Figure 4a**). Similarly, mink-associated GWAS hits were more likely to be polymorphic within mink samples than other categories of iSNVs (**Figure 4b**). As expected, we also observe that deer-associated GWAS hits, and mink-associated GWAS hits are more frequently fixed (at 100% frequency) in their respective samples than other categories of iSNVs (**Supplementary Figure 9a-b**). By definition, GWAS hits for a particular animal are more frequently fixed in these species than in humans (Naderi et al., 2023). Moreover, there could be overlap in the samples used for GWAS and in our current analysis. However, the GWAS only used consensus sequences and did not consider within-host diversity. The enrichment of iSNVs in GWAS hits (**Figure 4**) is therefore not a trivial artifact of the dataset, and suggests that more GWAS hits than expected by chance are under selection within individual animals. While it is difficult to disentangle relaxed purifying selection from strong positive selection, it is clear that selective pressures are species-specific because both GWAS hits and repeated iSNVs are species-specific. The overlap between repeated iSNVs and GWAS hits is incomplete, suggesting that some iSNVs are adaptive within but not between hosts.

**Figure 4.**
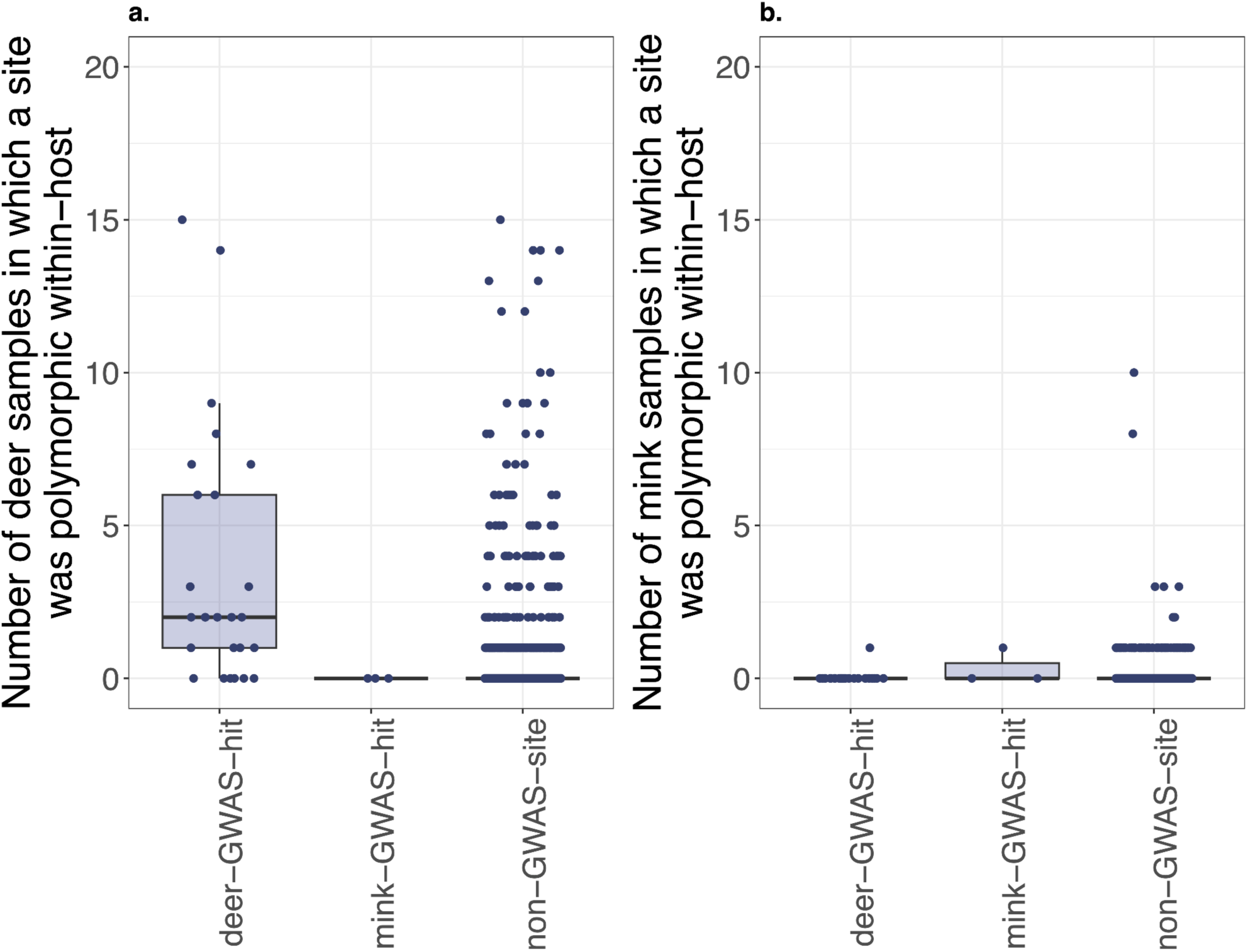
Deer- and mink-associated GWAS hits are more likely to be found as polymorphic (iSNVs) within hosts compared with other sites in the genome. The boxes display the interquartile range, with the midline marking the median, and whiskers extending up to 1.5x of the interquartile range, or data extremums. Datapoints beyond whiskers are outliers according to the interquartile criterion.

## Discussion

In our previous study, we identified 26 putative species-specific mutations in a sample of ∼100 SARS-CoV-2 consensus sequences sampled from deer, and only three in a sample of ∼1000 mink (Naderi et al., 2023). This suggests a greater amount of adaptive evolution in deer compared to mink, which could be explained by longer deer-to-deer transmission chains, as deer may be in less frequent contact with humans than farmed mink (Naderi et al., 2023, Lu et al., 2021). The high incidence of SARS-CoV-2 in deer – over 20% positivity rate in New York State – are also consistent with frequent deer-to-deer transmission (Feng et al., 2023). Longer deer-to-deer transmission chains, uninterrupted by passage through humans, could allow deer to accumulate deer-specific mutations that are rarely seen in humans.

Here, our analysis of within-host diversity across animal species revealed a higher level of polymorphism within individual deer when compared with humans. Why do deer harbor more within-host diversity than other animals? Possible explanations include higher mutation rate, longer infection duration, species-specific selective pressures, or higher rates of mixed-lineage infections in deer. Two measures of within-host diversity, the number of iSNVs and the pairwise nucleotide identity, π, were elevated in deer relative to humans, and these differences are not readily explained by mixed infections or technical confounders. Therefore, a combination of longer infections, higher mutation rates, and deer-specific selective pressures is likely to explain our observations. These possibilities are not mutually exclusive. For example, there is prior evidence for higher mutation rates in deer (McBride et al., 2023), which could explain the higher number of iSNVs. However, high mutation rate alone is unlikely to explain the enrichment of nonsynonymous mutations among repeated iSNVs in deer, which is a signature of natural selection.

Chronic infections of SARS-CoV-2 in immunocompromised humans have been extensively studied, indicating accelerated and potentially adaptive evolution within these patients (Borges et al., 2021). Many of the lineage-defining mutations that define globally successful SARS-CoV-2 variants of concern are also observed within chronic human infections, where they plausibly emerged (Harari et al., 2022). Similarly, chronic infections could be a viable explanation for the observed accelerated between-host evolution in deer (McBride et al., 2023) and for the higher within-host diversity observed in our data. Another possibility is that deer are no more likely to experience chronic infections than humans, but that they are systematically sampled at later stages of infection. This explanation is plausible if humans actively seek medical care (and associated sampling) shortly after (or even before) symptom onset, whereas deer are sampled opportunistically at random time points during infection. However, distinguishing between these possibilities is challenging given the available data.

Human chronic infections and deer infections with SARS-CoV-2 bear relevant similarities and differences. In both cases, viral evolution is likely adaptive, supported by the similarities of human chronic infection-derived mutations and those found in global variants of concern (Harari et al., 2022), and by the high number of GWAS hits in deer (Naderi et al., 2023). Here we showed that iSNVs in deer are enriched in deer-associated GWAS hits, suggesting that these mutations are under selection within hosts. We also observe an excess of nonsynonymous mutations among repeated mutations, another signature of positive selection. These repeated within-host mutations, as well the GWAS hits identified in a between-host analysis, are species-specific, suggesting species-specific selective pressures. This could involve a combination of species-specific positive selection or relaxation of purifying selection. The overlap between within-host mutations and between-host GWAS hits is statistically significant, but imperfect, indicating that some within-host mutations may be adaptive for onward transmission while others are not.

We have focused so far on deer, the species with the strongest evidence for species-specific adaptation and with the highest levels of within-host diversity, but we also made some other notable observations. First, we observed higher within-host diversity in lymph node tissue samples than nasopharyngeal samples in deer, though this observation was not statistically significant after controlling for confounding factors. It is biologically plausible for within-host diversity to be high in lymph tissue, as previous reports suggest compartmentalization of infection dynamics in different tissue types (Ke et al., 2022). However we cannot separate this from study-specific technical differences in our dataset, and our results on lymph node diversity therefore remain inconclusive. Second, mink, contrary to cats and dogs, were well-sampled and contained fewer mixed infections than humans. This could be explained by outbreaks of a single SARS-CoV-2 lineage, combined with early asymptomatic sampling during COVID-19 outbreaks on mink farms (Lu et al., 2021), limiting the potential for mixed-lineage infections. Third, we find that within-host diversity is not significantly different in experimental and natural infections. This result is counterintuitive, as we expect lower within-host diversity in experimental infections since these are inoculated with a single viral isolate, whereas natural infections can potentially be exposed to co-infection by different lineages. The effect of experimental infection is only interpretable for cats and dogs, as these are the only experimentally infected animals in the dataset, and we observe a higher median number of iSNVs in experimentally infected cats and dogs. All of our cat and dog samples were found to harbor only one lineage, which explains the lower Shannon diversity observed in these animals. Experimental cats and dogs were all sampled within 10 days of infection (Bashor et al., 2021), providing a rough benchmark for the expected amount of *de novo* mutation in that time frame. The observation that naturally infected cats and dogs harbor a similar number of iSNVs as in experimental infections suggests they were sampled in a similar time frame, all else being equal (*e.g.* viral loads and distinct selective pressures in experimental vs. natural settings). Unfortunately, we lack information about the duration of natural infections in our dataset.

It has been proposed that variants of concern such as Omicron could have arisen in a non-human animal host (Wei et al., 2021), but there remains no strong evidence that this has occurred to date. It is more plausible that a virus highly adapted to humans would arise from selection within a chronically-infected human host than within a different animal species, due in part to divergent species-specific selective pressures. On the other hand, it may be easier than previously thought for a population to evolve from one fitness peak (*e.g.* deer-adapted) to another (*e.g.* human-adapted) peak (Papkou et al., 2023). Viral evolution in non-human animals therefore merits continued surveillance because species-specific adaptation does not preclude future adaptation to humans. Together, our analysis identified deer as having higher within-host diversity than humans, providing a potential explanation for the rapid between-host evolution and adaptation of SARS-CoV-2 in deer. To confirm this explanation, we stress the need for further sampling of diverse animal species – and the analysis not just of consensus sequences, but of raw sequencing reads that can provide insight into within-host evolution.

## Acknowledgments

We are grateful to Sally Otto and Art Poon for providing constructive feedback on an earlier draft of the manuscript.

## Code availability

All code used for the analyses is available at https://github.com/Saannah/animalSARS-within-host

## Supplementary Tables

**Supplementary table 1:** Table of NCBI SRA identifiers for samples originally downloaded, and their respective metadata.

**Supplementary table 2:** Table of the metrics of interest for each sample, post quality control filtering.

**Supplementary table 3:** Table of the occurrence of previously identified GWAS hits in each sample.

**Supplementary table 4:** Table of coefficients for Model S1.

**Supplementary table 5:** Table of coefficients for Model S2.

**Supplementary table 6:** Table of coefficients for Model 2.

**Supplementary table 7:** Table of coefficients for Model S3.

**Supplementary table 8:** Table of samples with richness larger than one, and the breakdown of lineages called for each by Freyja.

**Supplementary table 9:** Table of coefficients for Model 4.

**Supplementary table 10:** Table of coefficients for Model S4.

**Supplementary table 11:** Table of coefficients for Model S5.

**Supplementary table 12:** Table of coefficients for Model S6.

**Supplementary table 13:** Table of repeated iSNVs observed in two or more deer (nasopharyngeal samples only), mink, or humans.

**Supplementary table 14:** Table of repeated iSNVs observed in two or more deer, including lymph node tissue samples.

## Supplementary Figures

**Supplementary figure 1.**
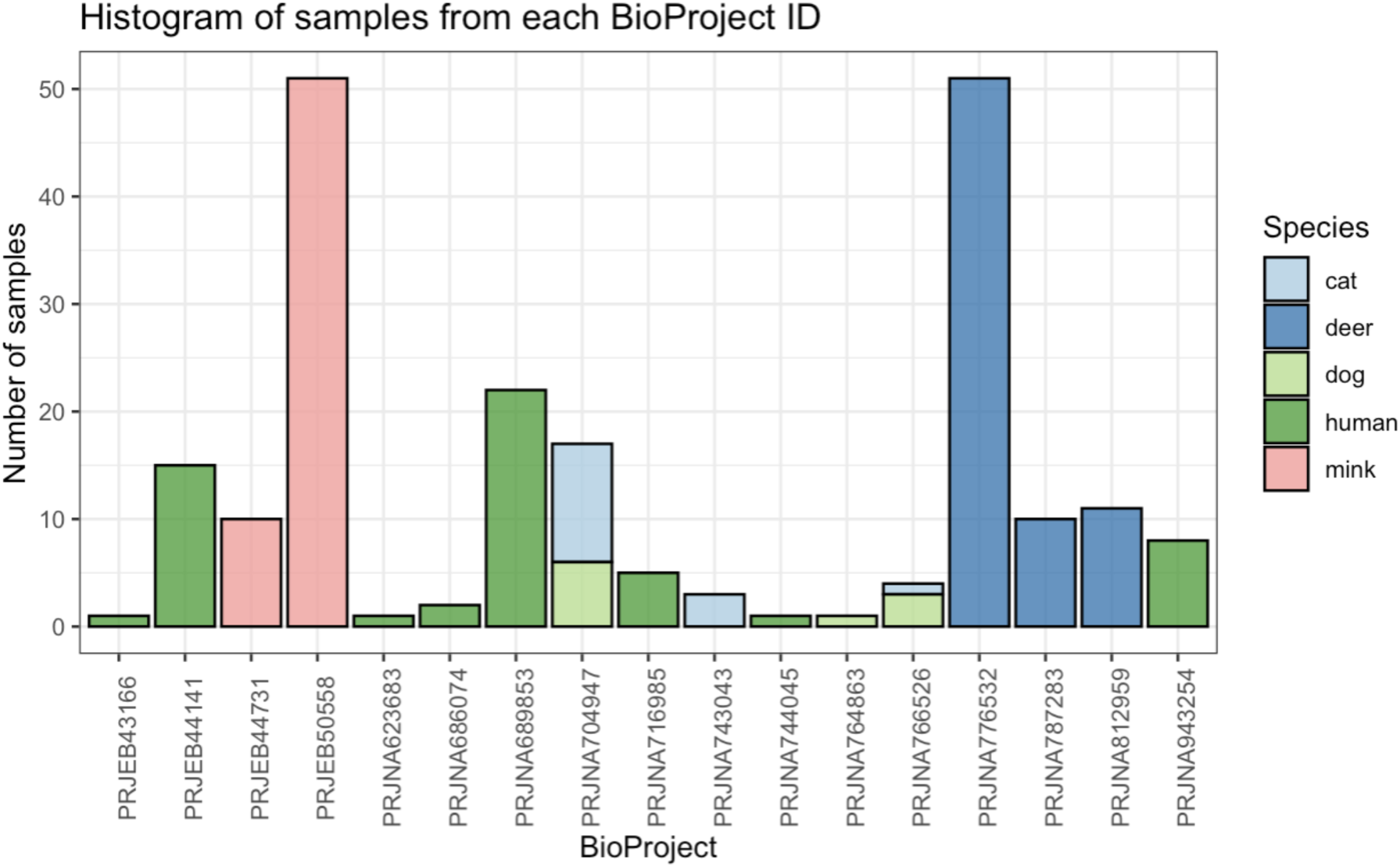
Histogram of the number of samples per BioProject ID, colored by species.

**Supplementary figure 2.**
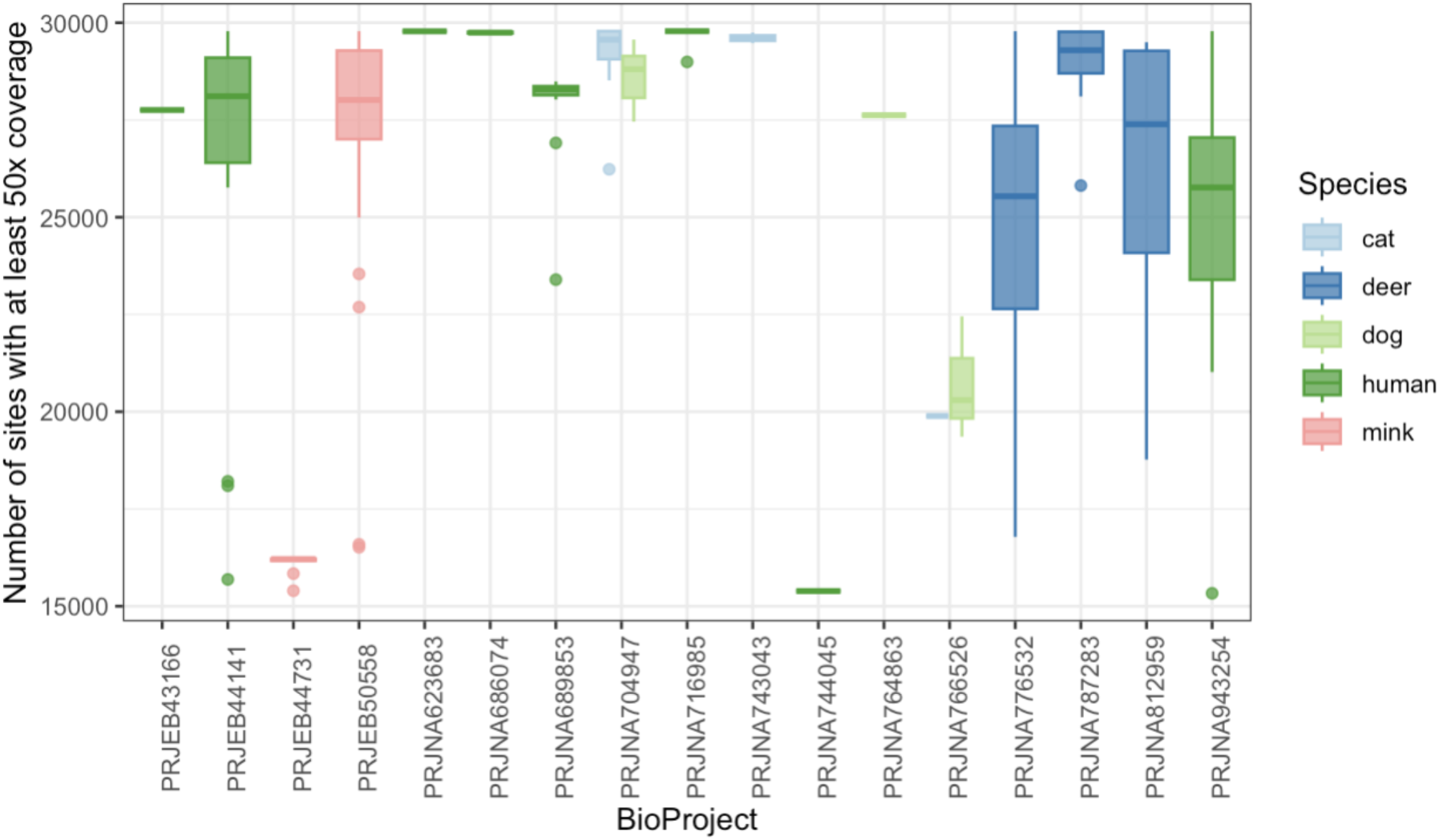
Distribution of the breadth of coverage in samples included in the analysis, broken down by BioProject ID and colored by species.

**Supplementary Figure 3.**
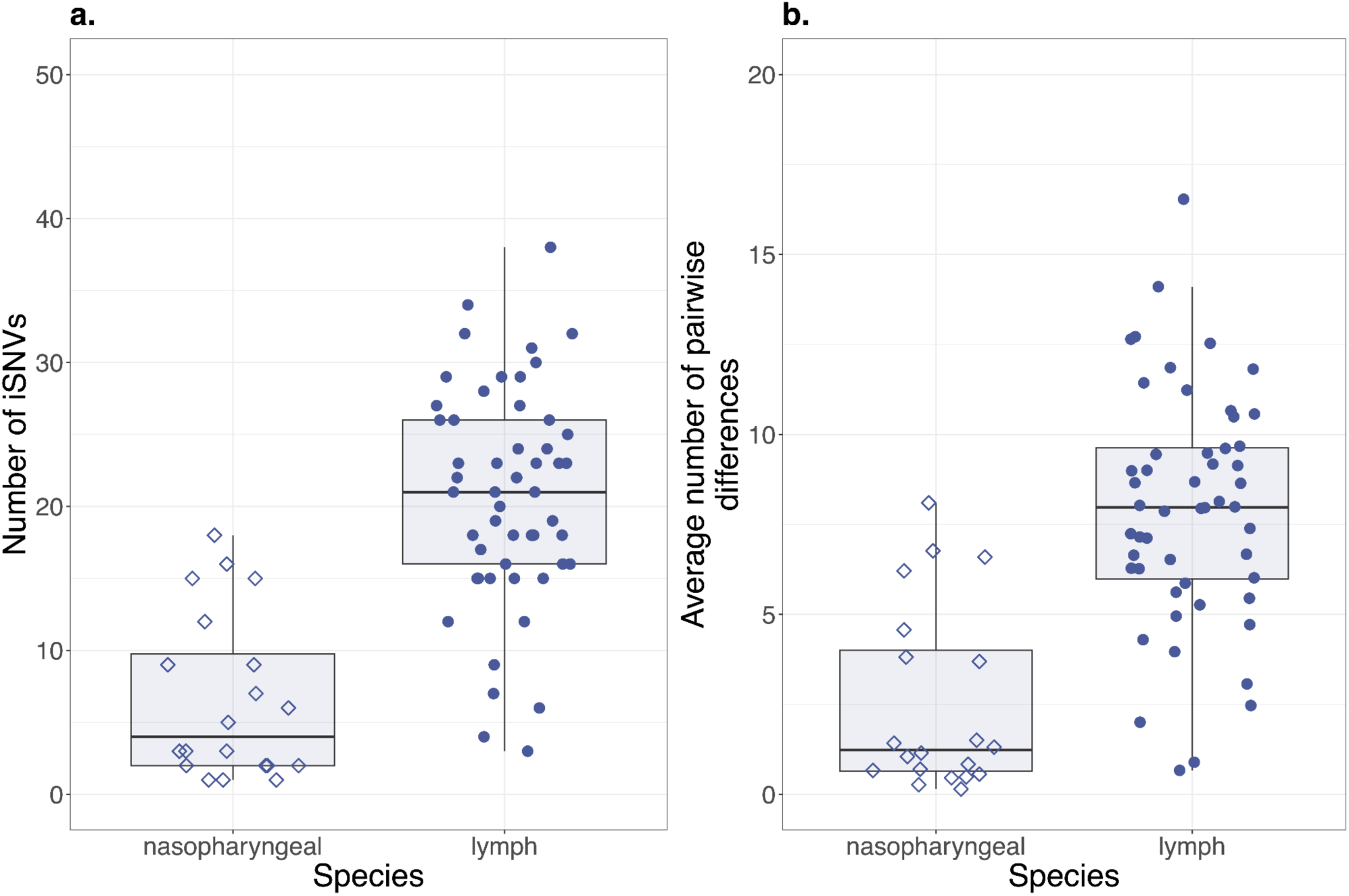
Within-host diversity in deer nasopharyngeal and lymph node samples, before controlling for possible confounders. Boxplots display (a) the number of iSNVs and, (b) the average number of pairwise differences (π) for deer across nasopharyngeal and lymph samples. Nasopharyngeal samples are shown with closed circles; lymph node tissue samples are shown with open diamonds. Raw data points are shown with random horizontal jitter. The boxes display the interquartile range, with the midline marking the median, and whiskers extending up to 1.5x of the interquartile range, or data extremums. Datapoints beyond whiskers are outliers according to the interquartile criterion.

**Supplementary figure 4.**
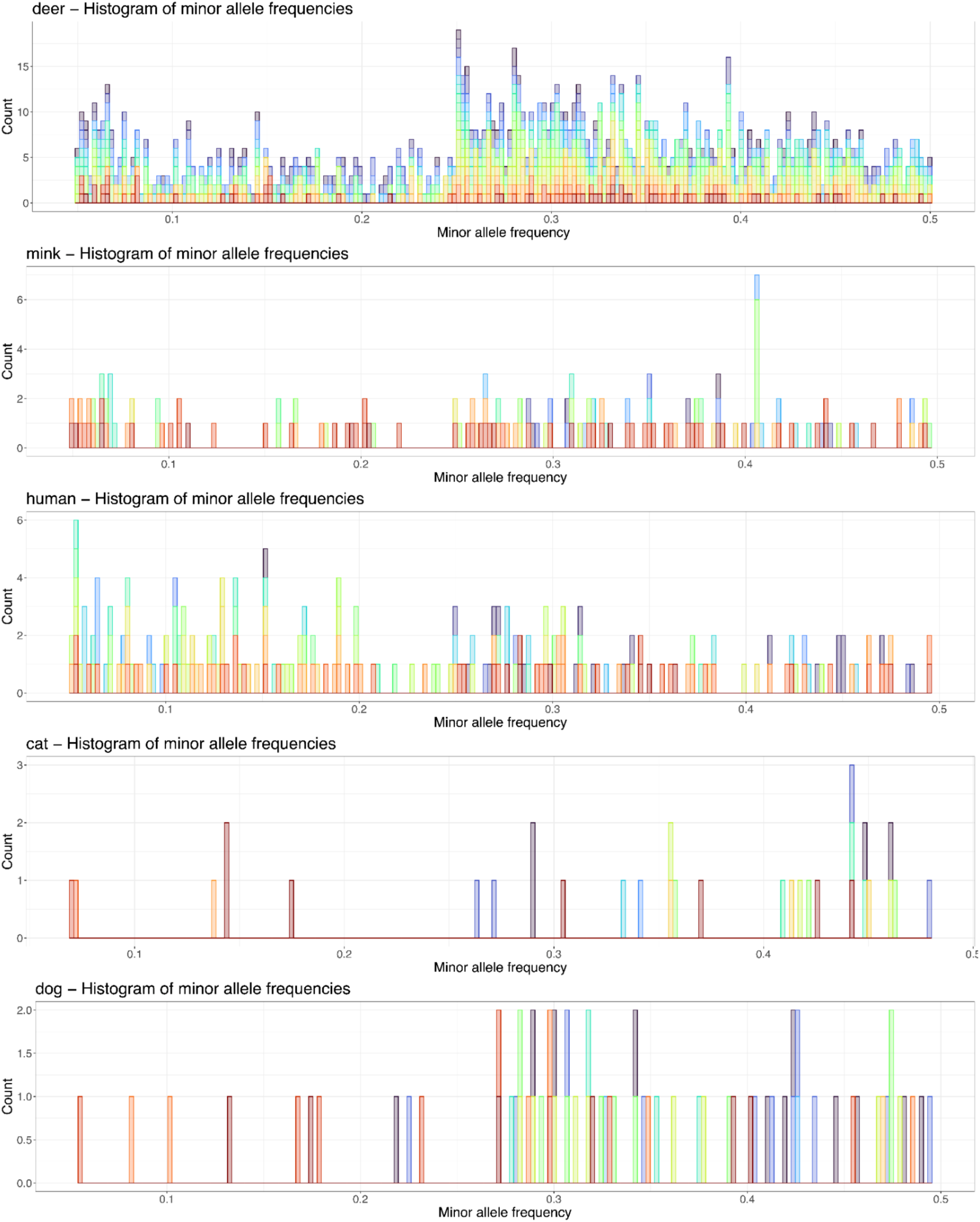
The distribution of minor allele frequencies of iSNVs found in different animal species. Each sample is shown in a unique color.

**Supplementary Figure 5.**
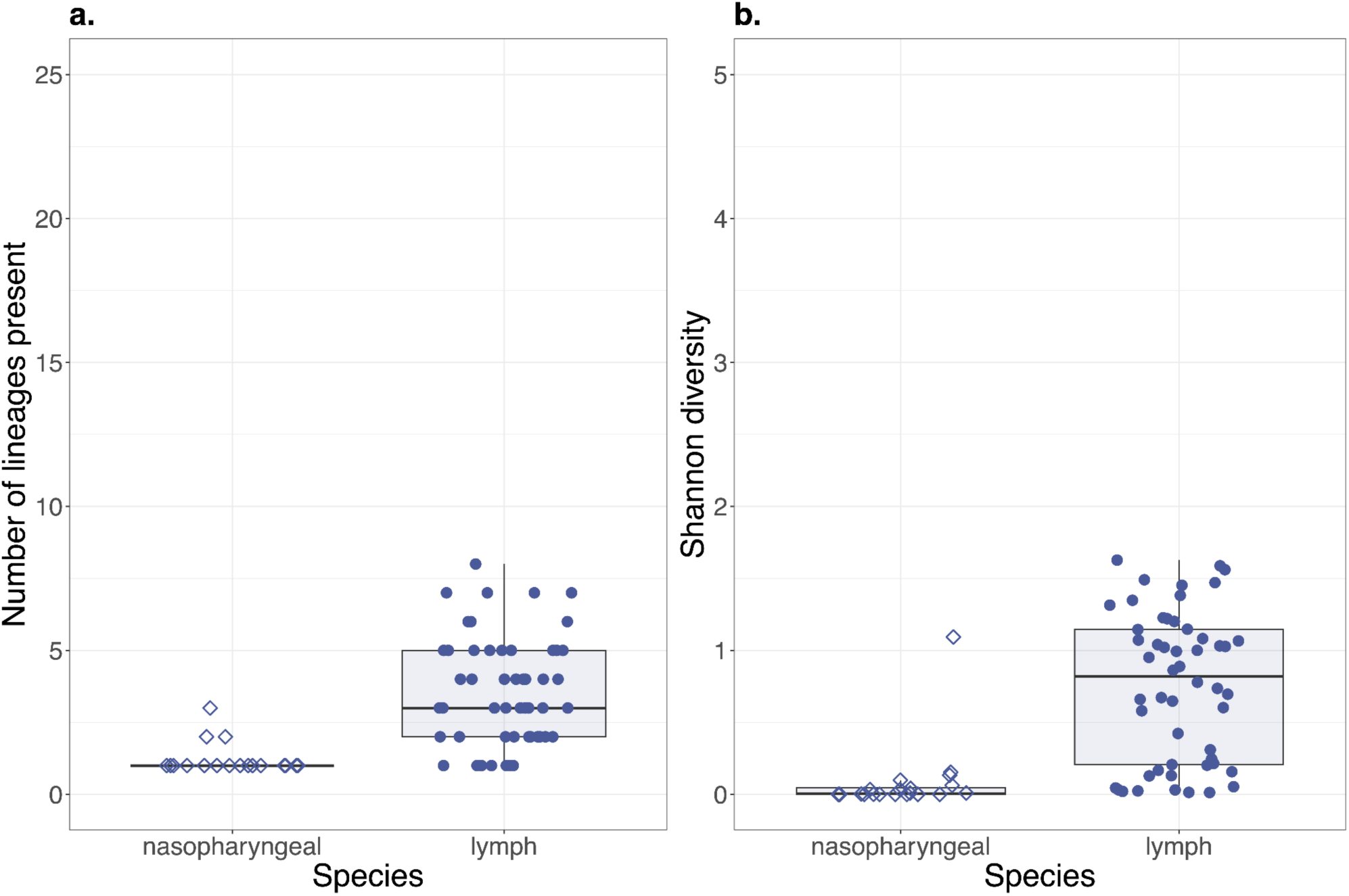
Higher median richness and Shannon diversity in lymph node samples. Boxplots display (a) the number of inferred lineages present and (b) the Shannon diversity index for deer across nasopharyngeal and lymph samples. Nasopharyngeal samples are shown with closed circles; lymph node tissue samples are shown with open diamonds. The boxes display the interquartile range, with the midline marking the median, and whiskers extending up to 1.5x of the interquartile range, or data extremums. Datapoints beyond whiskers are outliers according to the interquartile criterion.

**Supplementary Figure 6.**
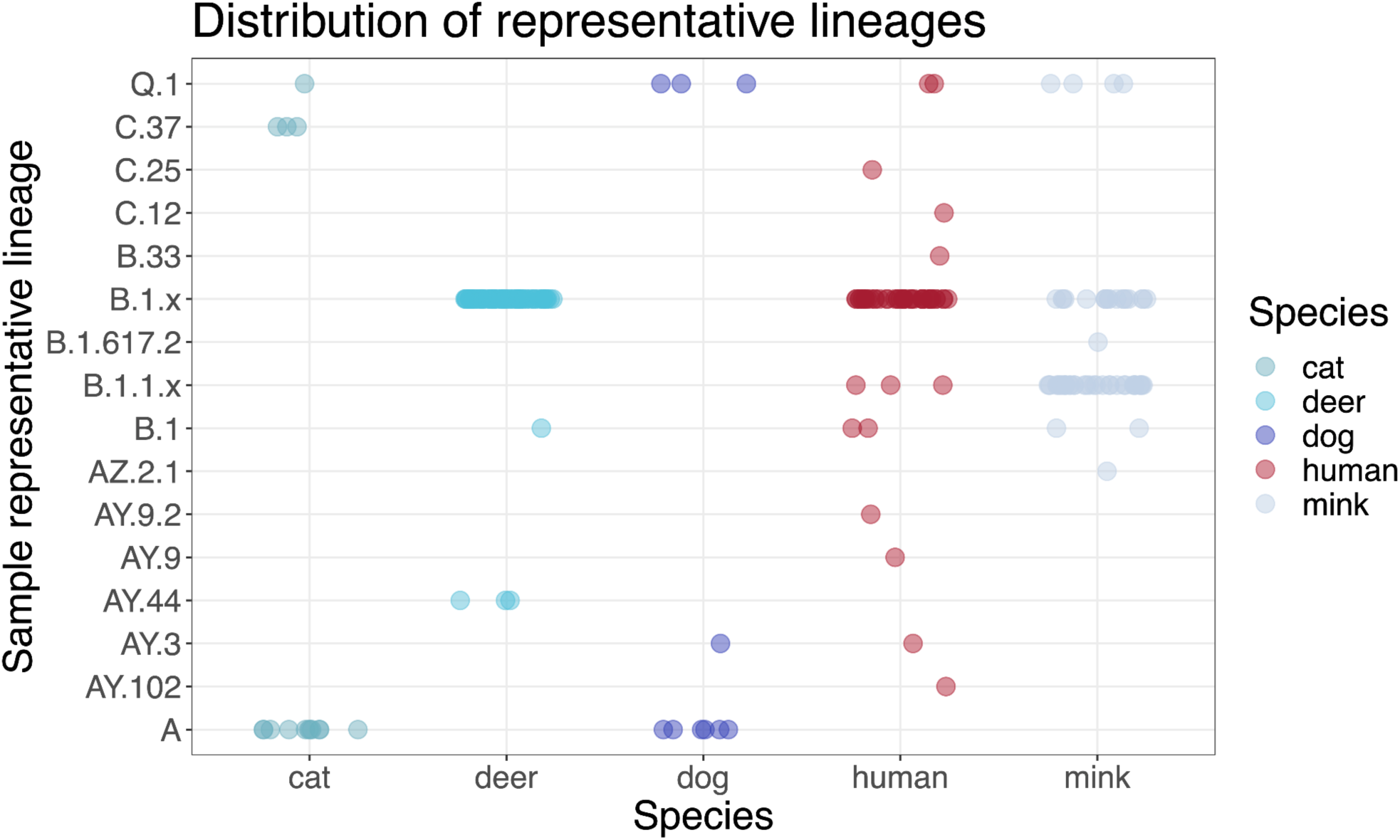
Distribution of representative (consensus) lineages across species. The major clusters of B.1.x and B.1.1.x in mink and deer are well-represented in their human matches. A clear discrepancy is the lab infected cats and dogs that are A lineages, and have no corresponding matches in human-host samples.

**Supplementary figure 7.**
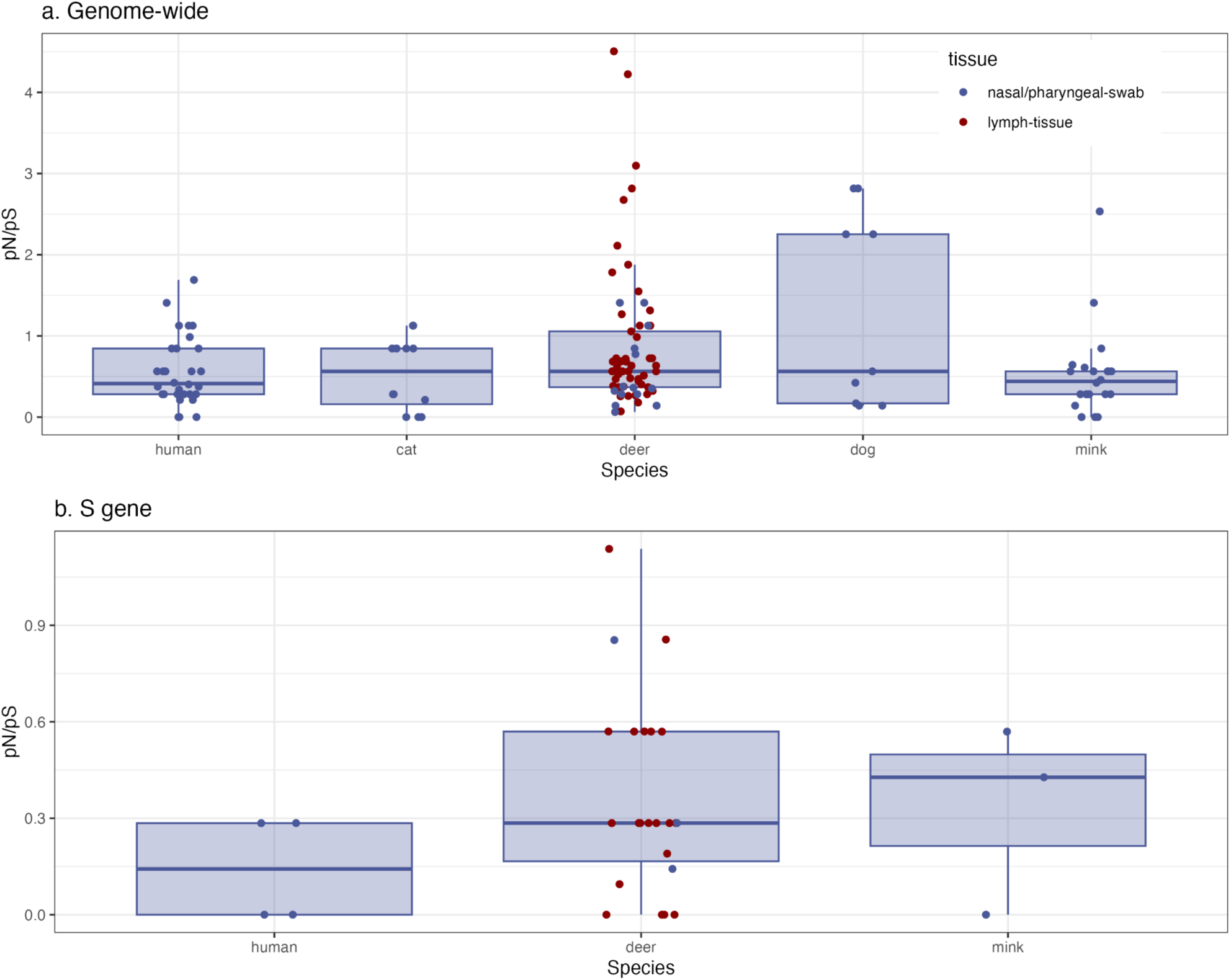
Scatter Plot of the pN/pS values calculated (A) genome wide and (B) in the S gene alone. Samples without any synonymous iSNVs were excluded from the analysis to avoid division by zero. As a result of this correction, all cat and dog samples were excluded in the analysis of the S gene. Samples from different tissue types (nasopharyngeal or lymph) are shown as blue or red points respectively.

**Supplementary figure 8.**
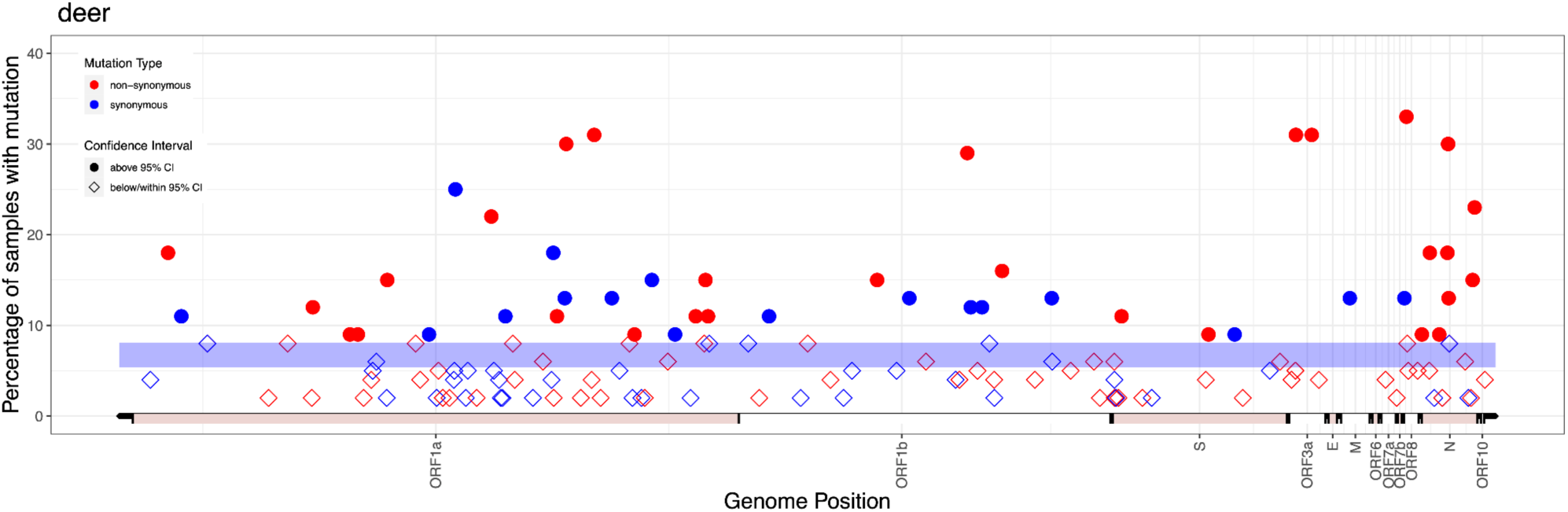
Repeated mutations in deer, with both nasopharyngeal and lymph node tissue samples included.

**Supplementary Figure 9.**
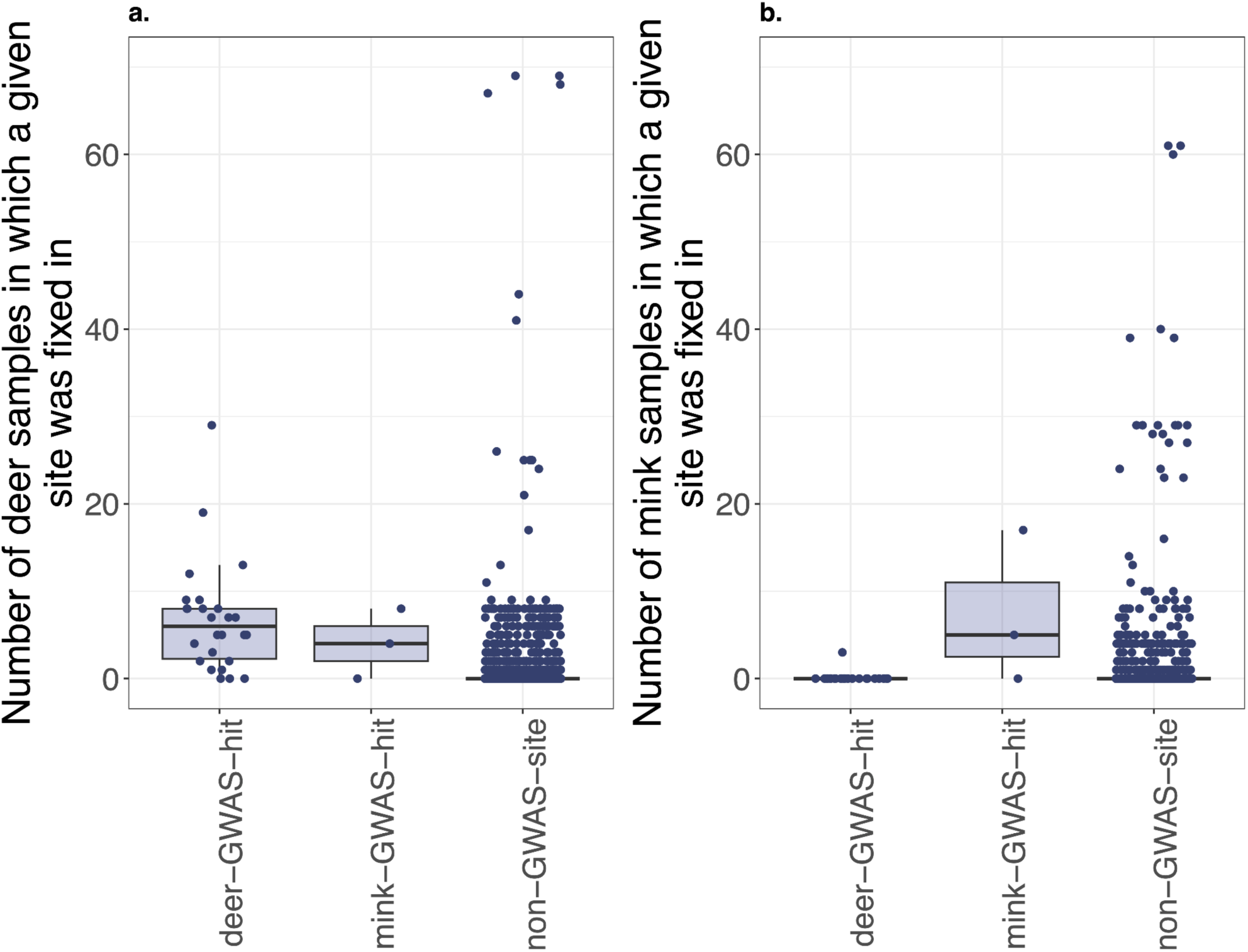
Deer- and mink-associated GWAS hits are more likely to be found as fixed SNVs in their respective host-species compared with other sites on the genome, confirming previous GWAS analyses. The boxes display the interquartile range, with the midline marking the median, and whiskers extending up to 1.5x of the interquartile range, or data extremums. Datapoints beyond whiskers are outliers according to the interquartile criterion.

## References

Andersen, K. G., Shapiro, B. J., Matranga, C. B., Sealfon, R., Lin, A. E., Moses, L. M., … & Sabeti, P. C. (2015). Clinical sequencing uncovers origins and evolution of Lassa virus. Cell, 162(4), 738–750.

Avanzato, V. A., Matson, M. J., Seifert, S. N., Pryce, R., Williamson, B. N., Anzick, S. L., … & Munster, V. J. (2020). Case study: prolonged infectious SARS-CoV-2 shedding from an asymptomatic immunocompromised individual with cancer. Cell, 183(7), 1901–1912.

Bashor, L., Gagne, R. B., Bosco-Lauth, A. M., Bowen, R. A., Stenglein, M., & VandeWoude, S. (2021). SARS-CoV-2 evolution in animals suggests mechanisms for rapid variant selection. Proceedings of the National Academy of Sciences, 118(44), e2105253118.

Bates, D., Mächler, M., Bolker, B., & Walker, S. (2015). Fitting Linear Mixed-Effects Models Using lme4. Journal of Statistical Software, 67(1), 1–48. 10.18637/jss.v067.i01

Borges, V., Isidro, J., Cunha, M., Cochicho, D., Martins, L., Banha, L., … & Gomes, J. P. (2021). Long-term evolution of SARS-CoV-2 in an immunocompromised patient with non-Hodgkin lymphoma. Msphere, 6(4), e00244–21.

Bourgey, M., Dali, R., Eveleigh, R., Chen, K. C., Letourneau, L., Fillon, J., … & Bourque, G. (2019). GenPipes: an open-source framework for distributed and scalable genomic analyses. Gigascience, 8(6), giz037.

Braun, K. M., Moreno, G. K., Wagner, C., Accola, M. A., Rehrauer, W. M., Baker, D. A., … & Moncla, L. H. (2021). Acute SARS-CoV-2 infections harbor limited within-host diversity and transmit via tight transmission bottlenecks. PLoS pathogens, 17(8), e1009849.

Choi, B., Choudhary, M. C., Regan, J., Sparks, J. A., Padera, R. F., Qiu, X., … & Li, J. Z. (2020). Persistence and evolution of SARS-CoV-2 in an immunocompromised host. New England Journal of Medicine, 383(23), 2291–2293.

Clark, S. A., Clark, L. E., Pan, J., Coscia, A., McKay, L. G., Shankar, S., … & Abraham, J. (2021). SARS-CoV-2 evolution in an immunocompromised host reveals shared neutralization escape mechanisms. Cell, 184(10), 2605–2617.

Feng, A., Bevins, S., Chandler, J., DeLiberto, T. J., Ghai, R., Lantz, K., … & Wan, X. F. (2023). Transmission of SARS-CoV-2 in free-ranging white-tailed deer in the United States. Nature Communications, 14(1), 4078.

Fox, J., & Monette, G. (1992). Generalized Collinearity Diagnostics. Journal of the American Statistical Association, 87(417), 178–183. 10.1080/01621459.1992.10475190

Fox J, Weisberg S (2019). An R Companion to Applied Regression, Third edition. Sage, Thousand Oaks CA. https://socialsciences.mcmaster.ca/jfox/Books/Companion/.

Garrison, E., & Marth, G. (2012). Haplotype-based variant detection from short-read sequencing. arXiv preprint arXiv:1207.3907.

Gonzalez-Reiche, A. S., Alshammary, H., Schaefer, S., Patel, G., Polanco, J., Carreño, J. M., … & van Bakel, H. (2023). Sequential intrahost evolution and onward transmission of SARS-CoV-2 variants. Nature Communications, 14(1), 3235.

Grubaugh, N. D., Gangavarapu, K., Quick, J., Matteson, N. L., De Jesus, J. G., Main, B. J., … & Andersen, K. G. (2019). An amplicon-based sequencing framework for accurately measuring intrahost virus diversity using PrimalSeq and iVar. Genome biology, 20, 1–19.

Harari, S., Tahor, M., Rutsinsky, N., Meijer, S., Miller, D., Henig, O., … & Stern, A. (2022). Drivers of adaptive evolution during chronic SARS-CoV-2 infections. Nature Medicine, 28(7), 1501–1508.

Karthikeyan, S., Levy, J. I., De Hoff, P., Humphrey, G., Birmingham, A., Jepsen, K., … & Knight, R. (2022). Wastewater sequencing reveals early cryptic SARS-CoV-2 variant transmission. Nature, 609(7925), 101–108.

Ke, R., Martinez, P. P., Smith, R. L., Gibson, L. L., Mirza, A., Conte, M., … & Brooke, C. B. (2022). Daily longitudinal sampling of SARS-CoV-2 infection reveals substantial heterogeneity in infectiousness. Nature Microbiology, 7(5), 640–652.

Kuznetsova A, Brockhoff PB, Christensen RHB (2017). “lmerTest Package: Tests in Linear Mixed Effects Models.” Journal of Statistical Software, 82(13), 1–26. doi:10.18637/jss.v082.i13.

Li, H., & Durbin, R. (2009). Fast and accurate short read alignment with Burrows–Wheeler transform. bioinformatics, 25(14), 1754–1760.

Lu, L., Sikkema, R. S., Velkers, F. C., Nieuwenhuijse, D. F., Fischer, E. A., Meijer, P. A., … & Koopmans, M. P. (2021). Adaptation, spread and transmission of SARS-CoV-2 in farmed minks and associated humans in the Netherlands. Nature Communications, 12(1), 6802.

Lythgoe, K. A., Hall, M., Ferretti, L., de Cesare, M., MacIntyre-Cockett, G., Trebes, A., … & Golubchik, T. (2021). SARS-CoV-2 within-host diversity and transmission. Science, 372(6539), eabg0821.

McBride, D. S., Garushyants, S. K., Franks, J., Magee, A. F., Overend, S. H., Huey, D., … & Bowman, A. S. (2023). Accelerated evolution of SARS-CoV-2 in free-ranging white-tailed deer. Nature Communications, 14(1), 5105.

Naderi, S., Chen, P. E., Murall, C. L., Poujol, R., Kraemer, S., Pickering, B. S., … & Shapiro, B. J. (2023). Zooanthroponotic transmission of SARS-CoV-2 and host-specific viral mutations revealed by genome-wide phylogenetic analysis. Elife, 12, e83685.

Nei, M., & Gojobori, T. (1986). Simple methods for estimating the numbers of synonymous and nonsynonymous nucleotide substitutions. Molecular Biology and Evolution, 3(5), 418–426.

NCBI: National Library of Medicine (US), National Center for Biotechnology Information; 2009 - [cited 2024/01/20]. Available from: https://www.ncbi.nlm.nih.gov/sra/

N’Guessan, A., Brito, I. L., Serohijos, A. W., & Shapiro, B. J. (2021). Mobile gene sequence evolution within individual human gut microbiomes is better explained by gene-specific than host-specific selective pressures. Genome Biology and Evolution, 13(8), evab142.

Papkou, A., Garcia-Pastor, L., Escudero, J. A., & Wagner, A. (2023). A rugged yet easily navigable fitness landscape. Science, 382(6673), eadh3860.

Pickering, B., Lung, O., Maguire, F., Kruczkiewicz, P., Kotwa, J. D., Buchanan, T., … & Bowman, J. (2022). Divergent SARS-CoV-2 variant emerges in white-tailed deer with deer-to-human transmission. Nature Microbiology, 7(12), 2011–2024.

R Core Team (2023). R: A Language and Environment for Statistical Computing. R Foundation for Statistical Computing, Vienna, Austria. <https://www.R-project.org/>.

Tan, C. C., Lam, S. D., Richard, D., Owen, C. J., Berchtold, D., Orengo, C., … & Balloux, F. (2022). Transmission of SARS-CoV-2 from humans to animals and potential host adaptation. Nature Communications, 13(1), 2988.

Tonkin-Hill, G., Martincorena, I., Amato, R., Lawson, A. R., Gerstung, M., Johnston, I., … & Wellcome Sanger Institute COVID-19 Surveillance Team. (2021). Patterns of within-host genetic diversity in SARS-CoV-2. Elife, 10, e66857.

Valesano, A. L., Rumfelt, K. E., Dimcheff, D. E., Blair, C. N., Fitzsimmons, W. J., Petrie, J. G., … & Lauring, A. S. (2021). Temporal dynamics of SARS-CoV-2 mutation accumulation within and across infected hosts. PLoS Pathogens, 17(4), e1009499.

Wei, C., Shan, K. J., Wang, W., Zhang, S., Huan, Q., & Qian, W. (2021). Evidence for a mouse origin of the SARS-CoV-2 Omicron variant. Journal of Genetics and Genomics, 48(12), 1111–1121.

Xue, K. S., & Bloom, J. D. (2020). Linking influenza virus evolution within and between human hosts. Virus Evolution, 6(1), veaa010.

Xue, K. S., Stevens-Ayers, T., Campbell, A. P., Englund, J. A., Pergam, S. A., Boeckh, M., & Bloom, J. D. (2017). Parallel evolution of influenza across multiple spatiotemporal scales. Elife, 6, e26875.

